# Mapping interactions of microbial metabolites and human receptors

**DOI:** 10.1101/614537

**Authors:** Dominic A. Colosimo, Jeffrey A. Kohn, Peter M. Luo, Sun M. Han, Amanda J. Pickard, Arka Rao, Justin R. Cross, Louis J. Cohen, Sean F. Brady

## Abstract

Despite evidence linking the human microbiome to health and disease, the mechanistic details of how the microbiota affects human physiology remain largely unknown. Metabolites encoded by bacteria are expected to play an integral role in the microbiota’s effect on its human host. Assigning function to these metabolites is therefore critical to determining the molecular underpinnings of the host-microbe relationship and ultimately developing microbiota inspired therapies. Here we use large-scale functional screening of small molecules produced by individual members of a simplified human microbiota to identify bacterial metabolites that agonize G-protein coupled receptors (GPCR). This analysis revealed a complex network of metabolite host receptor interactions and guided our identification of multiple microbiota derived agonists of GPCRs associated with diverse biological functions within the nervous and immune systems, among others. Collectively, the metabolite-receptor pairs we uncovered indicate that diverse aspects of human health are potentially modulated by structurally simple metabolites arising from primary bacterial metabolism.

**Statement of Significance:** Bacteria residing within the human body have been shown to influence human health. It is likely that physiological responses to the human microbiota are mediated by the collection of small molecules encoded within these bacteria. In this study we use direct functional screening of small molecules produced by individual members of a simplified human microbiota to identify new G protein coupled receptor-metabolite interactions that seek to explain the molecular underpinnings of the microbiota’s influence on its human host.

---

Human bodies are home to diverse and ever-changing collections of bacteria. The ability of the microbiota to influence human health has been explored extensively. [1] The most common methods for studying host-microbe interactions have featured “omics” based-analyses that have examined genomic, transcriptomic, proteomic or metabolic differences between patient cohorts. [2–5] Although these informatics-based methods have served as powerful tools for uncovering correlations between changes in the microbiota and health and disease, they are somewhat limited in their ability to reveal the mechanistic details of how the microbiota might alter mammalian physiology.[6] Much of the influence the microbiota has on its human host is likely encoded in the collection of small molecules it produces or modulates.[7] However, the number of well-defined interactions between metabolites produced by human associated bacteria and discrete human receptors is dwarfed by the number of reports attributing biological phenotypes to the microbiome, highlighting the need for a more systematic characterization of microbiota encoded bioactive metabolites.

In the case of synthetic small molecules that have proved useful for therapeutically modulating human physiology (*i.e.*, U.S. Food and Drug Administration (FDA) approved drugs) the majority (60-70%) function through just three classes of receptors: G-protein coupled receptors (GPCR), ion channels, or nuclear hormone receptors.[8] Many of these same proteins bind endogenous signaling molecules that regulate a wide range of physiological responses.[9] Based on the fact that these receptors play such an important role in how eukaryotic cells have evolved to translate external chemicals into biologic responses, it is likely the microbiota affects host physiology by modulating these same receptors with secreted metabolites.

Although healthy humans are colonized by hundreds, if not thousands of different bacterial species, the metabolic diversity they generate is likely limited by a high level of biosynthetic redundancy between bacterial species.[10] Due at least in part to this metabolic redundancy, it has been possible to use simplified human microbiomes (SIHUMIs) to model health and disease in murine models.[11, 12] In lieu of exploring random individual commensal species, we sought to conduct a more in-depth investigation of GPCR active microbiota encoded metabolites using bacteria from a model SIHUMI that contained a taxonomically diverse collection of commensal, health promoting and pathogenic bacteria. This consortium, which is composed of seven bacteria, assembled as a tool for studying gastrointestinal inflammation in the context of a healthy bacterial flora fulfills these general criteria and was therefore selected for use in this study.[13] Bacteria present in the SIHUMI consortium represent the major taxa found in the human microbiome and include beneficial bacteria (*Lactobacillus plantarum*, *Bifidobacterium longum*, and *Faecalibacterium prauznitzii*), non-pathogenic bacteria associated with disease (*Bacteroides vulgatus* and *Ruminococcus gnavus*), as well as clinically relevant pathogens (*Escherichia coli* LF-82 and *Enterococcus faecalis*).

We screened the metabolites produced by individually grown members of this SIHUMI consortium for agonism against 241 GPCRs. The resulting interaction map provides evidence, at the molecular level, for the existence of a complex network of microbial-host interactions, many of which involve receptors that have been modulated therapeutically with synthetic small molecules. Our characterization of interactions predicted by this analysis led to the discovery of both previously unrecognized as well as known microbiota encoded GPCR agonists. The structures of the active molecules we identified support the growing notion that simple bacterial metabolites arising from primary metabolic processes are likely to broadly impact human physiology.

## Results

### Culturing bacteria and GPCR screening

Bacteria from the SIHUMI consortium were individually fermented under anaerobic conditions in separate large scale (20L) culture vessels (Figure 1). After 10 days of static fermentation at 37 °C, hydrophobic resin was added directly to each culture. The resulting suspension was mixed to allow organic metabolites present in the fermentation broth to bind to the absorbent resin. Metabolite loaded resin was then collected by filtration, washed, and the bound metabolites were eluted with acetone. Each resulting crude metabolite extract was partitioned into 9 metabolite-rich fractions using reversed phased flash chromatography. A small aliquot of each fraction was arrayed for use in high-throughput GPCR screening. The remaining material was saved for follow up assays and for use in molecule isolation and structure elucidation studies. Although this pre-fractionation process increases the number of samples to be screened it simplifies the complexity of the crude culture broth extracts, which improves the signal in the primary screen thereby increasing the diversity of interactions that are identified and facilitating the downstream isolation of bioactive compounds. In addition to the bacterial fermentations, media not inoculated with bacteria was processed under identical conditions to control for possible bioactivity of small molecules derived directly from the media. The resulting library of bacterial metabolites was then screened with a cell-based assay for fractions that could agonize members of a panel of 241 GPCRs (Table S1). Specifically, a collection of recombinant cell lines engineered to measure β-arrestin recruitment by individual GPCR targets (β-arrestin recruitment assay) was used. For GPCRs with well characterized endogenous ligands, a maximum value for β-arrestin recruitment (100%) was set by exposing the recombinant cell line to a known agonist (Table S1). In the case of orphan receptors (*i.e.*, receptors without well characterized endogenous ligands), normalization of β-arrestin recruitment was performed by assigning the vehicle control to 0% activity. Hits were classified as such if a fraction induced a GPCR response to >30% of the control ligand (>50% for orphan GPCRs) and the comparable media control fraction showed <30% activity against the same GPCR (<50% for orphan GPCRs).

**Figure 1.**
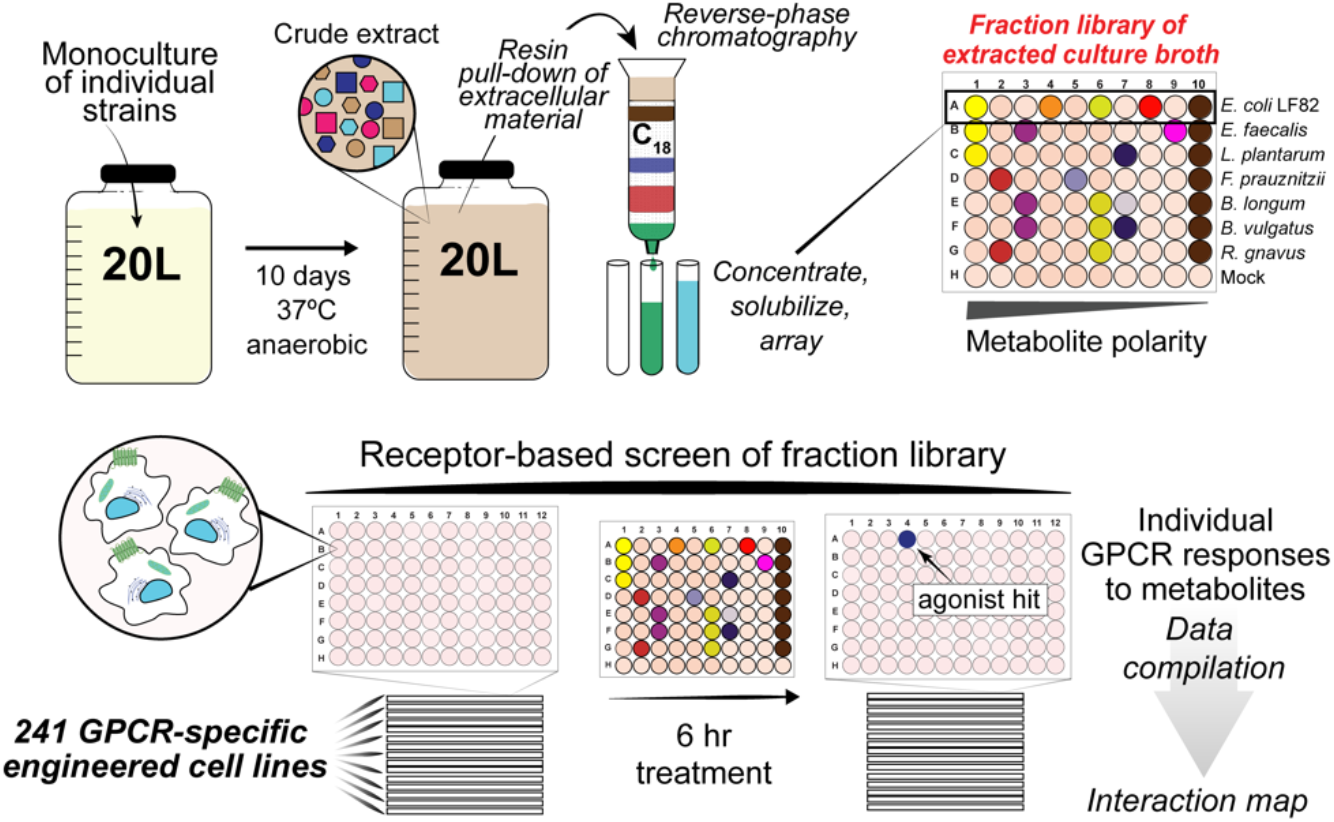
Experimental procedure for generating and screening library of secreted bacterial metabolites from large-scale monocultures of SIHUMI consortium members. This library was screened for the ability to agonize 241 distinct GPCRs.

The bacterial fraction library induced β-arrestin recruitment above our hit threshold levels for 67 of the 241 individual GPCR reporter cell lines we tested (Figure 2A-B, Table S2-S3). Of these 67 GPCRs, 54 did not show a strong background signal from the corresponding media control fraction, suggesting they were responding to bacterially encoded metabolites. Manual review of these 54 hits led us to de-prioritize 15 of these GPCR-fraction pairs due to high background of either the receptor or fraction (Table S4). The remaining 39 GPCRs were re-assayed in replicate; 22 of these GPCRs showed reproducible β-arrestin recruitment in response to 1 or more bacterial fractions (Figure 2C). Based on data from the Human Protein Atlas most of the receptors that reproducibly responded to bacterial metabolites are expressed at body sites regularly exposed to the microbiota (Figure 2C).[14] Notably, a large number of the receptors that were reproducibly agonized by microbiota encoded metabolites are also targeted by FDA approved drugs, indicating that receptors with proven physiological relevance are potentially modulated by bacterial ligands (Figure 2C). To identify specific GPCR-active metabolites, we used bioassay-guided isolation to purify metabolites from the large-scale culture broth fractions and *de novo* structure elucidation methods to determine their structures.

**Figure 2.**
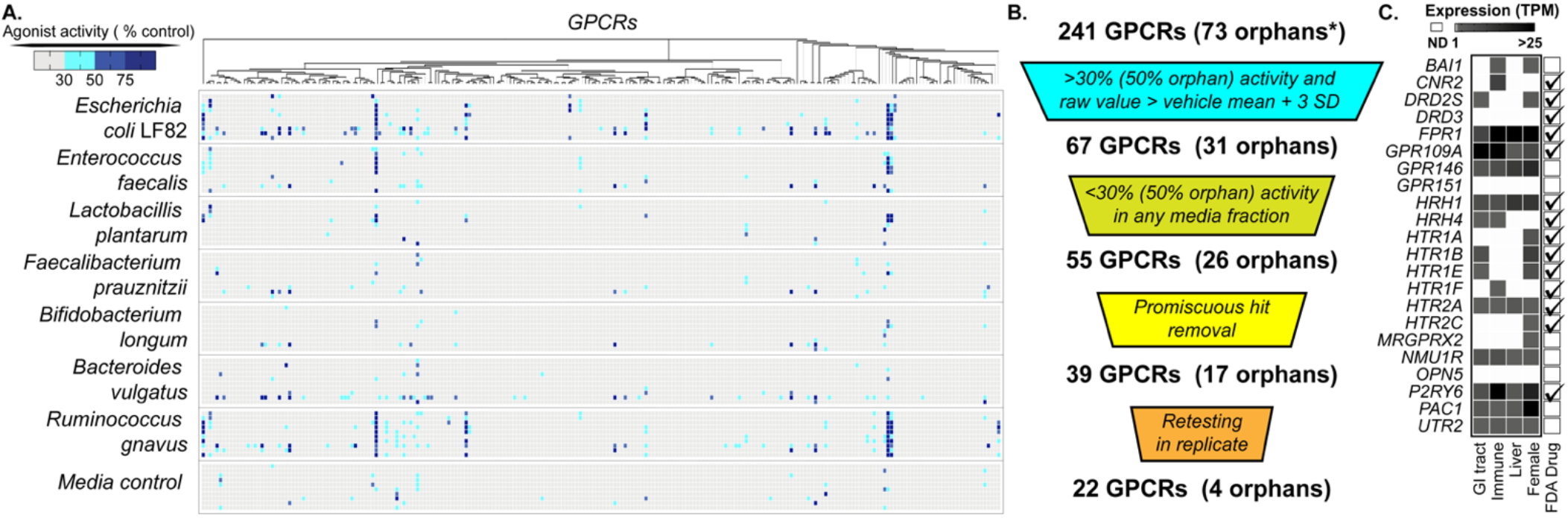
Overview of GPCR screening results. **A)** Heat map of individual assays for each GPCR tested indicating β-arrestin recruitment response normalized to endogenous or synthetic control compound (100%). **B)** GPCR hit prioritization scheme. **C)** Subset of GPCRs that show <30% (50% for orphans) response to the media control, but have >30% response (50% for orphans) to a bacterial fraction. The orphan receptors in this pool are BAI1, GPR146, GPR151 and OPN5. Receptor gene expression levels in tissues commonly exposed to the human microbiome (TPM, transcripts per million). Data is from the Human Protein Atlas.[14]. Receptors targeted by approved FDA drugs are indicated on the right. [52]

### Known and novel ligands for hydroxycarboxylic acid receptors

A number of receptors agonized in our screen are known to respond to bacterial ligands. As an initial validation exercise we characterized activities expected to arise from well-known bacterial GPCR agonists. The hydroxycarboxylic acid receptors, GPR81, GPR109A and GPR109B, are agonized by both human and bacterial ligands.[15] Bioassay guided fractionation of GPR109A active fractions from cultures of both *L. plantarum* and *R. gnavus* yielded nicotinic acid (Vitamin B3) as the active metabolite (Figure 3A, Figure S1). Nicotinic acid, an essential nutrient acquired either through diet or gut bacteria, is the most extensively studied non-endogenous ligand for this receptor. Its ability to regulate lipid metabolism in hyperlipidemic patients is well established in the clinic.[16] The identification of this well characterized and *in vivo* validated ligand receptor pair suggests that data generated in our screen has the potential to uncover new biologically relevant metabolite GPCR interactions.

**Figure 3.**
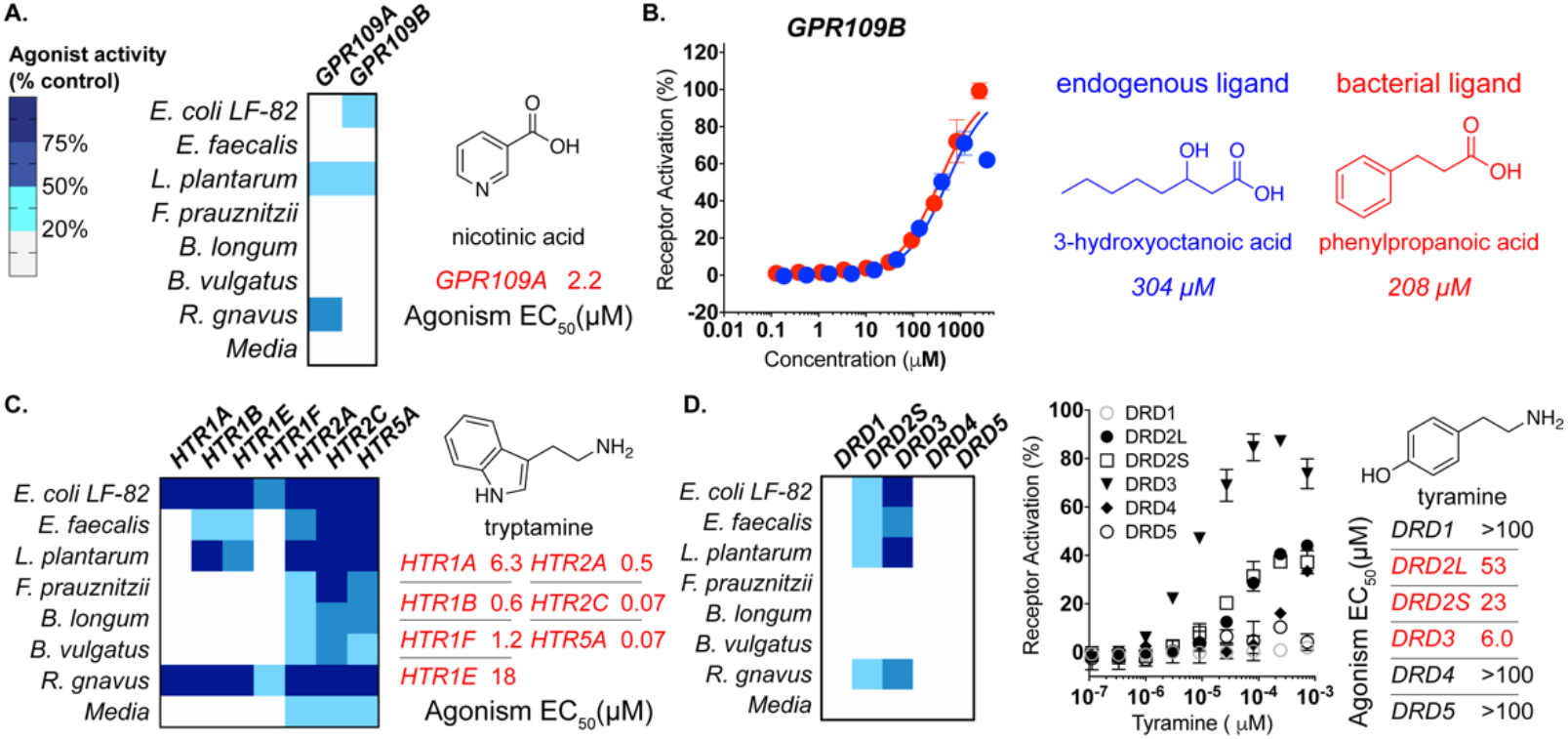
Bacterial ligands for hydroxycarboxylic acid and neurotransmitter receptors. The single fraction with maximum activity for each bacterial strain is depicted in heat maps. **A)** Left: Heat map depicting agonism of GPR109A and GPR109B by bacterial fractions. Right: Agonist activity (EC_50_) of purified nicotinic acid against GPR109A. **B)** Left: Dose response curves (DRCs) for known and novel GPR109B agonists (right). **C)** Left: Heat map depicting agonism of HTR receptors by culture broth extract fractions. Right: Agonist activity (EC_50_) of tryptamine against HTRs. **D)** Left: Heat map depicting agonism of DRD family receptors by culture broth extract fractions. Right: Agonist activity (EC_50_) of tyramine against DRDs. All dose-response curves were run in duplicate. Error bars are standard deviation. Error bars that are shorter than height of the symbol are not shown.

Fractions derived from cultures of both *E. coli LF82* and *L. plantarum* agonized a second hydroxycarboxylic acid receptor, GPR109B. Bioassay guided fractionation did not identify the endogenous ligand produced in humans, 3-hydroxyoctanoic acid, but instead yielded phenylpropanoic acid as the active metabolite (Figure 3B, Figure S1). This previously unknown GPR109B agonist, elicited a similar GPCR response than did 3-hydroxyoctanoic acid (Figure 3B). While the EC_50_ values for the known and novel ligands (304 μM and 208 μM, respectively) are higher than is often seen for endogenous GPCR ligands (Table S1), no more potent GPR109B agonists have been identified, outside of those derived synthetically.[17] Whether this is an inherent attribute of the receptor or represents a failure to identify the natural human ligand for this receptor remains to be seen.

Phenylpropanoic acid is not produced natively by humans. Its presence in human fecal and sera samples has been attributed to either *de novo* biosynthesis by bacteria or microbial transformation of dietary compounds, most notably by species of *Clostridium*.[18–21] While we believe phenylpropanoic acid is the first microbiota derived agonist to be identified for GPR109B, in a screen of synthetic molecules, several aromatic D-amino acids were found to be GPR109B agonists that can trigger chemoattraction signaling pathways in leukocytes.[22, 23] In a quantitative analysis of human fecal water, phenylpropanoic acid was reported in healthy patients at an average concentration of 77.30 μg/mL (513 μM).[18] At this concentration, production of phenylpropanoic acid by gut bacteria would be high enough to expect agonism of GPR109B. An analysis of the concentration of phenylpropanoic acid in a larger number of patient samples will be required to determine whether phenylpropanoic acid dependent agonism of GPR109B is likely to be a common phenomenon in the gut.

### Known and novel ligands for neurotransmitter receptors

Our GPCR interaction map revealed numerous bacterial fractions that strongly agonized neurotransmitter receptors, a key component of the gut brain axis (Figure 1C).[24] Bacterial produced aromatic amines, most notably tryptamine, have recently been reported as agonists of neurotransmitter receptors, particularly serotonergic GPCRs (5-hydroxytryptamine receptors, HTRs).[25] A majority of bacteria in this SIHUMI produced fractions that agonized HTRs (Figure 3C). Isolation of the active metabolite yielded tryptamine, which was produced in varying quantities by members of this SIHUMI (Figure S3). These results agree with various reports that HTRs are responsive to a wide array of bacteria due to the generality of tryptamine production across species.[26]

In fractions from multiple bacterial species we observed agonism of the D2-type dopamine receptors (DRDs), DRD2 and DRD3 (Figure 3D). Bioassay-guided isolation led to the aromatic amine tyramine as the major metabolite responsible for DRD agonism in these fractions. Tyramine arises from decarboxylation of tyrosine and differs from dopamine only by the absence of a second hydroxyl on the aromatic ring. It is reported to accumulate to μM levels in the gastrointestinal tract, a phenomenon which has been attributed to production by human microbiota.[27] While no biological significance has been assigned to the microbiota dependent accumulation of tyramine in animal models, it is sufficiently potent that its observed concentration in the gastrointestinal tract is high enough to agonize D2 subtype DRDs.

In contrast to the broad activation seen for DRDs and HTRs across extracts from all of the bacteria in this consortium, a specific response to fractions from *E. coli* LF82 was detected for a member of the histamine receptor (HRH) family, HRH4. Our inability to retain HRH4-activity when using hydrophobic chromatography during the bioassay guided purification process suggested that the active molecule was highly polar. We did not, however, expect that the activity was due to bacterially produced histamine, as the active fraction did not agonize other HRH family receptors and we could not detect histamine by LC-MS or NMR. We ultimately found the polyamine cadaverine to be the metabolite responsible for HRH4 agonism (Figure 4A). The activity of cadaverine was confirmed using a commercial standard (EC_50_ 1.1 μM) (Figure 4B). In addition to cadaverine, bacteria commonly produce a number of other simple polyamines including agmatine, spermidine and putrescine.[28] To explore the promiscuity of HRH4 agonism by polyamines, we tested synthetic standards of these metabolites for the ability to induce β-arrestin recruitment by each member of the HRH receptor family. Agmatine and putrescine showed limited activity against HRH4 (Figure 4C), while spermidine did not show activity against any receptor in the family. The inability for humans to biosynthesize cadaverine suggests that the influence of polyamines on histamine signaling pathways is likely specific to bacterial metabolism.

**Figure 4.**
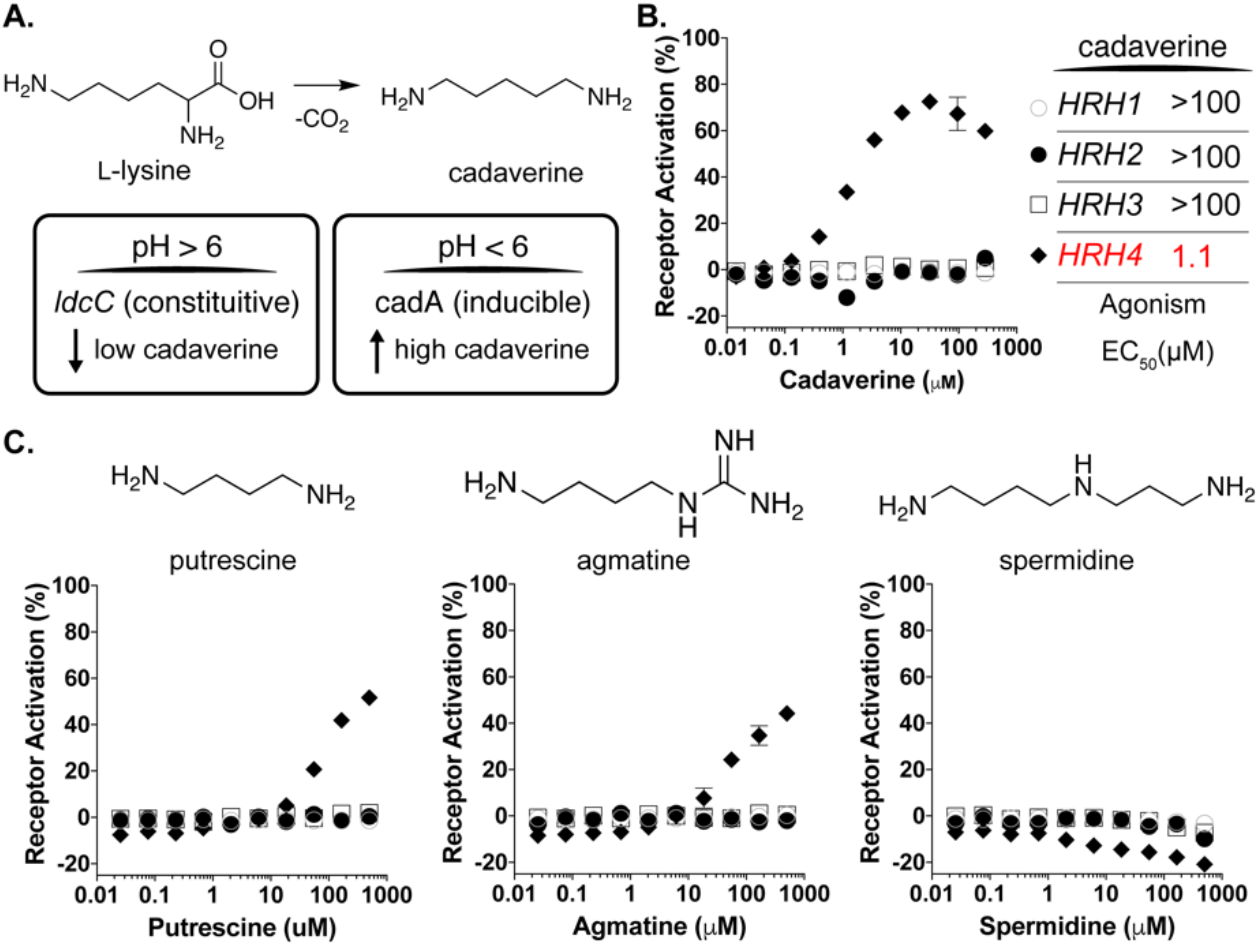
Cadaverine is a bacterial ligand for a specific histamine receptor. **A)** Top: Schematic of cadaverine biosynthesis from L-lysine. Bottom: Bacterial enzymes that catalyze this reaction include LydC, which is constitutively expressed and CadA, whose gene expression is induced at low pH. **B)** Dose response curves for cadaverine against HRH family receptors. **C)** Dose response curves (bottom) of bacterial polyamines (above) against HRH family receptors. Receptor symbols are labeled as in panel B. All dose-response curves were run in duplicate. Error bars are standard deviation. Error bars that are shorter than height of the symbol are not shown.

Cadaverine is produced by bacteria through the decarboxylation of lysine (Figure 4A), while agmatine and putrescine are derived from arginine. In a number of bacteria, including many associated with the microbiota (Table S5), cadaverine is encoded by both the constitutive *ldc* gene cluster as well as the *cad* gene cluster, which is induced at low pH (pH <6.8).[29] High level production of cadaverine by the CadA lysine decarboxylase plays a role in protecting against acid stress.[30]. As the pH of the digestive system varies longitudinally and features multiple acidic sections (*e.g.*, cecum pH ~5.7), high level production of polyamines by *cad* gene cluster containing bacteria is likely to occur at numerous sites in the GI tract. The biological relevance of gastrointestinal production of polyamines remains unclear, however host responses to polyamines have been reported in various contexts.[31, 32] Interestingly, although histamine receptor subtypes differ in their associated functions and their distribution throughout the human body, HRH4 is expressed in the gastrointestinal tract and altered expression levels have been linked to inflammatory responses that are related to inflammatory bowel diseases and cancer.[33]

A growing number of studies have uncovered connections between gut microbiota and the nervous system.[34, 35] Our exploration of microbiota encoded neurotransmitter receptor agonists expands the mechanistic evidence for simple biogenic amines serving as potentially widespread modulators of the gut-brain axis.[14] These data imply that microbiota-dependent dopaminergic, serotonergic and histaminergic responses likely represent general signaling events in the gastrointestinal tract with varying activation profiles depending on the specific collection of bacteria present in an individual’s microbiome.

### Structurally distinct lipids agonize diverse GPCRs

Lipids, which represent diverse GPCR active ligands [36, 37], predominantly elute very late in our fractionation protocol (Figure 5A). Based on the receptor interaction map we could initially classify GPCRs as lipid responsive if they were agonized by the late lipid-enriched fractions of the extract library. A subset of receptors, including GPR120, CNR2, GPR171, GPR132, responded broadly to the lipid fraction from most of the consortium, whereas other responses were specific to particular species (BAI1, NMU1R, UTR2). HPLC-charged aerosol detection analysis of the lipid fractions indicated they contained not only mixtures of simple saturated fatty acids but also other more complex lipid species (Figure 5B). Marrying unique receptor activity profiles with unique lipid signals guided us to previously unrecognized bacteria encoded GPCR agonists.

**Figure 5.**
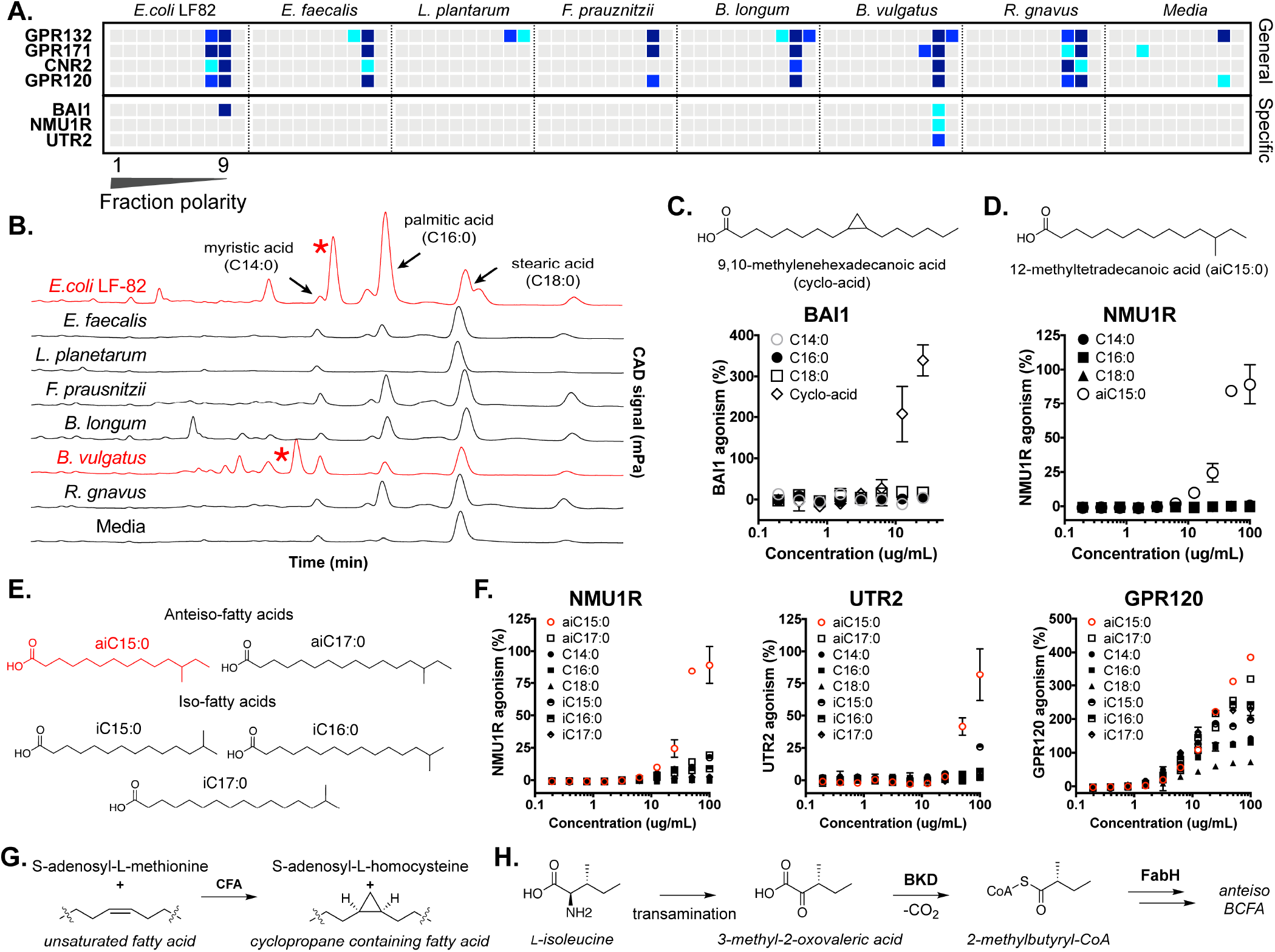
Lipid responsive GPCRs. **A)** Heat map of GPCRs demonstrating general (top) or specific (bottom) responses to lipid-rich fractions of bacterial extracts. **B)** Overlaid CAD chromatograms with common lipids and unique lipids (red asterisk) marked. **C)** Structure of BAI1-active lipid 9,10-methylenehexadecanoic acid isolated from *E. coli* LF82 and response of BAI1 to various fatty acids **D)** Structure of NMU1R-active lipid, 12-methyltetradecanoic acid, isolated from *B. vulgatus* and response of NMU1R GPCR to various fatty acids. **E)** Panel of branched chain fatty acids tested for GPCR fidelity. **F)** Response of NMU1R, UTR2 (specific) and GPR120 (general) to branched chain fatty acid panel. **G)** Biosynthesis of cyclopropane rings from unsaturated fatty acids using cyclopropane fatty acid synthase (CFA). **H)** Early steps in the biosynthetic scheme for ante-iso branched chain fatty acids (BCFAs) in bacteria (BKD, branched-chain α-keto acid dehydrogenase; FabH, β-ketoacyl-acyl carrier protein synthase III). All dose-response curves were run in duplicate. Error bars are standard deviation. Error bars that are shorter than height of the symbol are not shown.

The brain angiogenesis factor 1 (BAI1) receptor was agonized by lipid fractions from the Gram-negative bacteria in the consortium: *E. coli* and *B. vulgatus*. The *E. coli* LF82 lipid fraction showed the most potent agonism of BAI1 and therefore it was selected for further analysis. Bioassay guided fractionation identified the BAI1 agonist as the cyclopropyl-containing lipid 9,10-methylenehexadecanoic acid (EC_50_ 11 μM). Synthetic 9,10-methylenehexadecanoic acid, but no saturated lipids we tested agonized BAI1, confirming the specificity of the receptor reflected in the initial GPCR activity map (Figure 5C). The enzyme cyclopropane-fatty-acyl-phospholipid synthase (Cfa) uses the one carbon donor S-adenosyl-L-methionine to generate cyclopropyl lipids from unsaturated fatty acids (Figure 5G). Cyclopropane-containing fatty acids are important membrane components in Gram-negative as well as mycolic acid bacteria.[38] Macrophages use BAI1 as a pattern recognition receptor to sense Gram-negative bacteria and to induce selective phagocytosis and antimicrobial responses; 9,10-methylenehexadecanoic acid may represent a previously unrecognized recognition motif for innate immune responses.[39–42]

Two peptide receptors NMU1R (neuromedin receptor 1), which mediates satiety and peristalsis in the gut [43, 44] and the vasoconstriction inducing urotensin 2 receptor (UTR2) responded specifically to lipid fractions generated from *B. vulgatus*. Isolation of the active metabolite yielded the *anteiso*-methyl branched-chain fatty acid 12-methyltetradecanoic acid (aiC15:0) (Figure 5D). *Anteiso*-fatty acids (ai) contain an alkyl branch at the *ante*-penultimate carbon in contrast to iso-fatty acids (i) which branch at the penultimate carbon. Both synthetic and natural aiC15:0, but no simple fatty acids we tested, agonized NMU1R (EC_50_ 125 μM) and UTR2 (EC_50_ 191 μM). Lipid sensitivity of NMU1R and UTR2 appears specific to aiC15:0, as fatty acids with even slightly modified branching patterns (iC15:0) or carbon chain length (aiC17:0) displayed minimal agonist activity (Figure 5E-F). Methyl branched fatty acids arise from the use of a branched primer in place of acetyl CoA in normal fatty acid biosynthesis. In the case of *anteiso*-methyl-branched fatty acids, 2-methyl-butyryl-CoA, which is derived from isoleucine is used to prime fatty acid biosynthesis (Figure 5G). The selectivity for branched primers lies with the β-ketoacyl acyl carrier protein synthase (KAS III or FABH) that carries out the first condensation in fatty acid biosynthesis. *Anteiso*-methyl fatty acids are predominantly produced by Gram-positive FABH enzymes.[45, 46] Roughly 10% of bacteria have lipid pools enriched in branched chain fatty acids.[45] *B. vulgatus*, is among those bacteria enriched in branched chain fatty acids and maintains aiC15:0 as ~30% of its total fatty acid repertoire.[47]

Bacteria are known to produce diverse and oftentimes taxa specific, collections of lipids. The examples described here from examining even this minimized model microbiome suggest the potential for markedly different receptor activation profiles and hence biological consequences depending on the specific lipid signature encoded by an individual’s microbiome. For BAI1, NMU1R and UTR2 our data suggests that they differentially respond to lipids produced by largely Gram-positive or Gram-negative bacteria indicating that their activities will fluctuate with changes in the gross taxonomic composition of a microbiome.

### Analysis of mice colonized with the seven strain SIHUMI consortium

In parallel with our *in vitro* screen studies, we used high-resolution mass spectrometry-based metabolomics to compare germ free and SIHUMI consortium colonized mice. For this analysis, cultures of individually grown bacteria from the consortium were combined and the mixed sample was gavaged into germ-free C57BL/6 mice. PCR based species analysis of DNA extracted from the stool of animals three days post inoculation confirmed their colonization by the consortium. Ten days post colonization the lumen material (cecal stool) was collected from the germ-free controls as well as the SIHUMI colonized animals. Using targeted high-resolution mass spectrometry, we looked for differences in metabolite accumulation in these samples (Figure 6).

**Figure 6.**
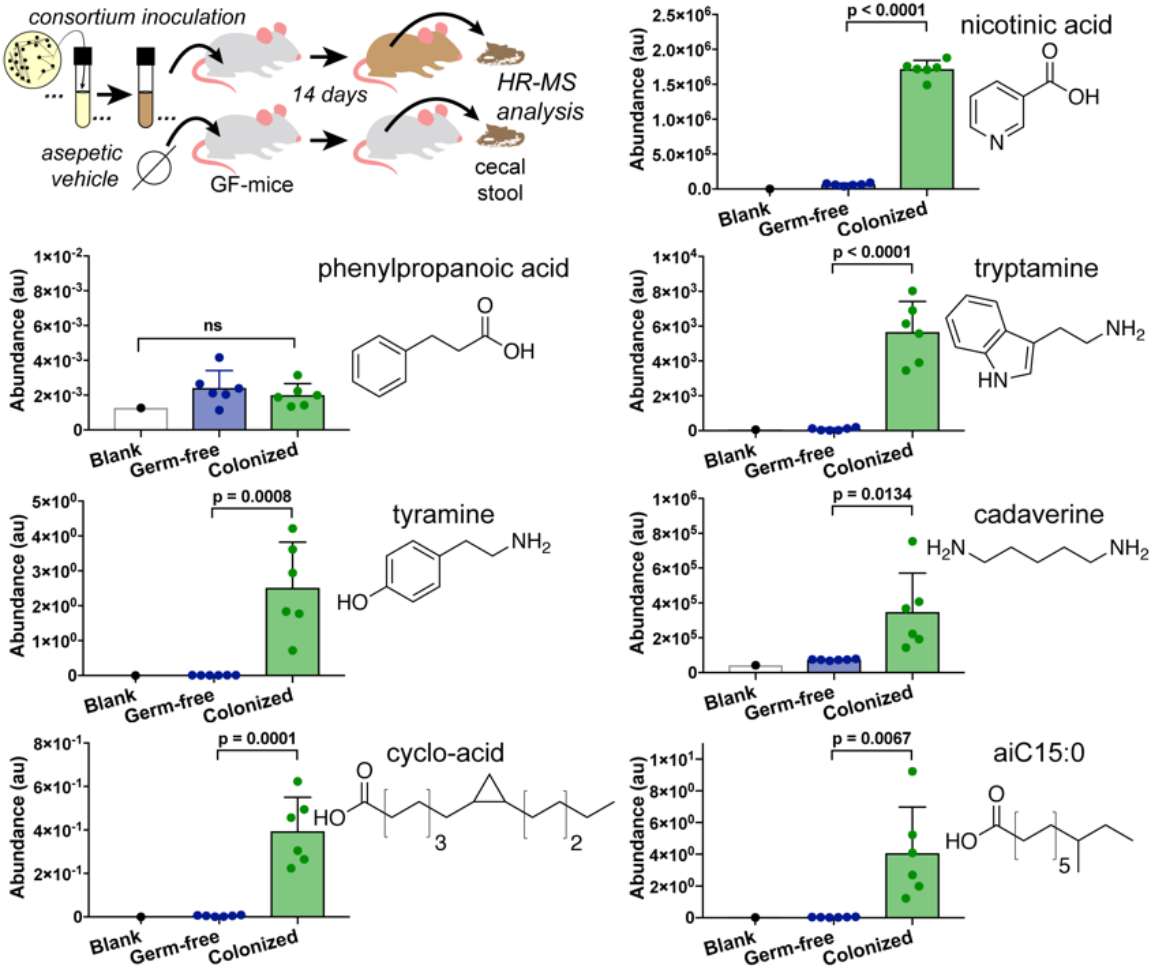
Comparative analysis of metabolite levels in the cecum of abiotic mice to levels in mice inoculated with SIHUMI consortium. Metabolite presence in lumen cecal samples was determined by high-resolution mass spectrometry (see Methods for experimental details). Samples were normalized to each other based on the addition of isotopically labeled internal standards during extraction, n=6 mice (3 female and 3 male), error bars are standard deviation, p values are derived from the unpaired t test.

Targeted MS analysis of cecum extracts revealed that all but one of the GPCR-active metabolites we identified was enriched in these mice compared to their abiotic counterparts (Figure 6, Table S7), suggesting a largely parallel biosynthesis in laboratory grown mono-cultures and the consortium *in vivo*. The lone exception was phenylpropanoic acid. Our failure to identify phenylpropanoic acid in this specific model study does not preclude its potential biologic relevance in other settings, especially in light of the fact that it has been seen at μM levels in human patient samples.[18] The lack of *in vivo* production is likely due to low production of this metabolite by the specific strain used in this consortium, which is supported by the low GPR109B activity we observed in our original fraction screen.

## Discussion

Phenylpropanoic acid, cadaverine, 9-10-methylenehexadecanoic acid, and 12-methyltetradecanoic acid add to a growing list of structurally simple molecules that are capable of modulating human signaling pathways that underlie diverse clinically relevant areas of physiology including immune recognition, neurotransmission, and inflammation.[35, 48] The biosynthetic simplicity of these metabolites combined with their abundant starting materials likely drives their high titers in the gut and potential broad biological relevance. Expanding functional screening to include not only more bacteria but also additional culture conditions and receptor families will undoubtedly provide additional insight into the biochemical mechanisms and small molecules underlying human-microbiome interactions. For example, a study that was published while the work reported here was under review examined a different collection of bacteria and identified different GPCR active metabolites.[49] Advancements in laboratory culturing techniques now allow for a majority of gut bacteria to be cultured from fecal samples.[50, 51] Systematic functional screening of metabolites produced by this growing collection of bacteria is likely to be a rewarding avenue for developing mechanistic hypotheses that can be tested in specific animal models.

## Materials and Methods

Full description of the experimental procedures used can be found in Supplemental Material.

## Acknowledgements

All bacterial strains were generously provided by Daniel Mucida. High-resolution mass spectrometry of purified compounds was performed by Rockefeller University Proteomics Core. We are grateful to C. Fermin, E. Vazquez, and G. Escano in the Precision Immunology Institute at the Icahn School of Medicine at Mount Sinai (PrIISM) Gnotobiotic facility and Microbiome Translational Center for their help with gnotobiotic experiments. Funding was provided by the Bill and Melinda Gates Foundation (OPP1168674) and the National Institutes of Health (5R01AT009562–02).

## Author Contributions

D.A.C., L.J.C. and S.F.B. designed experiments. D.A.C., J.A.K. and P.M.L. performed analytical chemistry. S.M.H. and L.J.C. performed murine experiments. A.J.P., A.R. and J.R.C. designed, performed, and analyzed mass-spectrometry experiments. D.A.C. and S.F.B. wrote manuscript.

## Materials and Methods

### Media construction

LBM media: LBM media was derivative of a media recipe previously utilized in our laboratory. For 1 L media: Bring 17 g/L brain heart infusion, 5 g/L yeast extract, 200 mg MgSO_4_•7H_2_O, 100 mg MnCl2•4H_2_O up in 800 mL deionized water and autoclave for 30 minutes liquid cycle. After coming to room temp, add supplements (final concentrations: 5 μg/L hemin, 1 g/L maltose, 1 g/L cellobiose, and 0.5g/L L-cysteine), which can be made ahead of time and stored, protected from light, at −20 °C in aliquots, except hemin which can be stored at 4 °C and L-cysteine which should be made fresh. Use autoclaved deionized water to bring final volume to 1 L. For culturing anaerobes: place media in anaerobic chambers for at least 48 hrs to allow diffusion with anaerobic gas.

### Cultivation of bacteria

Bacterial strains of the SIHUMI consortium used listed in Table S2. Cultures of <1 L: Anaerobic bacteria were cultured in an incubator set to 37 °C placed inside of vinyl anaerobic chamber (Coy) with a gas mix of 5% CO_2_, 5% H_2_, and 90% N_2_. Cultures of >1 L: When cultivating bacteria for construction of bacterial extract library, bacteria were inoculated in 1 L or 2 L media bottles (Chemglass) inside anaerobic chamber, then sealed with anaerobic septa (Chemglass) and moved into large walk-in 37 °C incubator constructed from a 6×6×12 light protective tent (APOLLO^®^ HORTICULTURE) outfitted with a regulator (INKBIRD^®^), heat source (VORNADO^®^), and ventilation system (IPOWER^®^). Freezer stocks of the SIHUMI cohort were generously donated by the Mucida laboratory (Rockefeller University). Freezer stocks were thawed and bacteria were cultivated overnight in LBM media until turbid. These bacteria were streaked onto LBM agar plates and upon growth, single colonies were picked, cultivated overnight, and genotyped (GeneWiz). Upon confirmation of genetic identity, these same cultures were used to generate colony 20% glycerol stocks that would be used for the entirety of the study. Bacteria specific primers were designed to allow for PCR-based identification of each specific strain. *Escherichia coli* LF-82 (F: GTTAATACCTTTGCTCATTGA, R: ACCAGGGTATATAATCCTGTT, 340 bp); *Faecalibacterium prausnitzii* DSM 17677 (F: CCCTTCAGTGCCGCAGT, R: GTCGCAGGATGTCAAGAC, 158 bp); *Enterococcus faecalis* OG1RF (F: CCCTTATTGTTAGTTGCCATCATT, R: ACTCGTTGTACTTCCCATTGT, 144 bp);*Bifidobacterium longum* ATCC 15707 (F: GGGTGGTAATGCCGGATG, R: TAAGCGATGGACTTTCACACC, 442 bp); *Lactobacillus plantarum* ATCC BAA-793 (F: AGCAGTAGGGAATCTTCCA R: CACCGCTACACATGGAG, 341 bp); *Bacteroides vulgatus* ATCC 8482 (F: GGTGTCGGCTTAAGTGCCAT, R: CGGAYGTAAGGGCCGTGC, 140 bp); *Ruminococcus gnavus* ATCC 29149 (F: CGGTACCTGACTAAGAAGC, R: AGTTTYATTCTTGCGAACG, 429 bp) For large scale fermentations the following protocol was used: Bacterial stocks were thawed and used to inoculate 5 mL LBM liquid cultures that were cultivated overnight. The next day, species specific primers were used to confirm identity (as described below) and upon passing purity check, these 5 mL cultures were used to inoculate 500 mL LBM at a ~1:100 ratio. After turbidity was reached, an aliquot of the 500 mL culture was removed and PCR was performed with universal 16s rRNA primers 27F (AGAGTTTGATCMTGGCTCAG) and 1492R (GGTTACCTTGTTACGACTT). The CR product was subject to Sanger sequencing (GeneWiz) and upon passing inspection for the correct species, the 500 mL culture was used to inoculate 12 L of LBM media at a 1:100 inoculation ratio. The 20 L cultures were cultivated, protected from light at 37 °C, for 10 days without shaking. Amerlite XAD-7HP (Sigma Aldrich) was aliquoted in 20 g increments and activated by soaking in methanol for 10 minutes, followed by 5 washes with deionized water to remove excess methanol. After 10 days, activated Amberlite XAD-7HP was added to the cultures (20 g dry weight/L) and the slurries were gently shaken (90 rpm) on a tabletop shaker for 4 hrs. After incubation with the cultures, the resin was removed via cheese-cloth filtration and the collected resin. alongside the cheese-cloth, was placed inside a 1 L Fernbach flask to which 1.5 L acetone was added. This acetone elution was allowed to occur for 2 hrs with shaking (150 rpm), after which the organic solvent was collected and fresh acetone, of equivalent volume, was added. This second elution was allowed to occur overnight with light shaking at 22 °C. Both elutions were added together and solvent was removed via rotary evaporation (Buchi) at 25 °C to afford the dry crude extract, which was stored at −20 °C until fractionated, as detailed below.

### Fractionation of bacterial extracts

Crude extracts (~1-3 g/12L) were re-suspended in ~300 mL methanol and the soluble material was decanted into a 500 mL round bottom flask (rbf). Free C_18_ resin (2-3 g) was added and the slurry was evaporated under reduced pressure using a rotary evaporator with temperature set to 25 °C (Buchi). The dry material was collected from the rbf, packed semi-tightly into a 50 g cartridge, and capped with a passive frit (Teledyne). This material was chromatographed over a 150 g C18 Gold (Teledyne ISCO) using a solvent system of water (Solvent A) and methanol (Solvent B), with no acid added, with the following conditions. 5 column volumes (CV) of 5% B, 5% B to 99% B over 10 CV, flush with 10 CV 99% B. All flow-through was collected in 50 mL tubes and combined as follows:

Solvent was evaporated using an SPD-2010 speedvac (Thermo Scientific) with a RH-19 rotor (Thermo Scientific) and the resulting dry material was weighed and resuspended at 100 mg/mL using ACS grade DMSO (Fisher Scientific). Of this solution, 250 µL was removed and added to 250 µL DMSO to create 500 µL 50 mg/mL solution; this solution was aliquoted into various sizes of 96-well plates for facile thawing and biological testing at a later time. The remaining 100 mg/mL solution was stored at −80 °C until validation studies required material for bio-assay guided fractionation.

### GPCR assays

GPCR activities were measured by Eurofins DiscoverX using the PathHunter^®^ β-Arrestin assay.[1] This assay uses β-galactosidase (β-Gal) that has been split into two inactive portions as a reporter to measure the activation of a GPCR. The β-Gal fragments are called EA for Enzyme Acceptor and ED for Enzyme Donor.(US Patent: US20090098588A1). Using these fragments, a unique reported cell line was created for each GPCR of interest. In each unique cell line the EA fragment is fused to β-Arrestin and the ED fragment is fused to the GPCR of interest. Upon GPCR activation, β-Arrestin recruitment to the receptor physically colocalizes the ED and EA fragments thereby restoring β-Gal activity. β-Gal complementation is measured using chemiluminescent PathHunter^®^ Detection Reagents. For our initial screen (Figure 2) all 80 culture broth extract fractions were screened in singleton against the Eurofins DiscoverX gpcrMAX (**168 GPCRs**, Table S2) and orphanMAX (73 GPCRs, Table S3) panels. All subsequent validation and bioassay guided fraction studies were run in at least duplicate using individual reporter cell lines for specific GPCRs of interest.

#### *Eurofins DiscoverX* generic agonist protocol

1. Sample is added to individual GPCR reporter cell lines grown in microtiter plates. Cells are incubated at 37 °C or room temperature for 90 or 180 minutes. 2. Assay signal is generated through addition of 12.5 or 15 μL (50% v/v) of PathHunter Detection reagent cocktail, followed by incubation for a one hour at room temperature. 3. Microplates are read with a PerkinElmer EnvisionTM instrument for chemiluminescent signal detection. 4. Compound activity is analyzed using the CBIS data analysis suite (ChemInnovation, CA). For receptors with known ligands, percentage activity is calculated using the following formula: % Activity =100% x (mean RLU of test sample — mean RLU of vehicle control) / (mean MAX control ligand — mean RLU of vehicle control). For orphan receptors, percentage inhibition is calculated using the following simplified formula: % Activity =100% x (mean RLU of test sample — mean RLU of vehicle control) / (mean RLU of vehicle control). For some orphan receptors that exhibit low basal signal, the more sensitive PathHunter Flash Kit is used.

### Cadaverine biosynthesis bioinformatics

Using the annotated genome of *E. coli* LF-82 the Pfam protein features of CadA (Accession ID LF82_0254) or the amino acid sequence itself were used as a query against the annotated genome collection provided by the NIH Human Microbiome Project.[2, 3] This dataset was chosen as it allowed us to confidently assign annotated bacterial genomes containing a *cadA* gene to body sites origin. For the Pfam analysis: three Pfam motifs are found within the CadA amino acid sequence: PF01276 constitutes the major domain of the decarboxylase, PF03711 constitutes the C-terminal domain and PF03709 constitutes the N-terminal domain. The raw data for both BlastP (>30% identity) and Pfam-based analyses are available in Table S5a and S5b, respectively.

### High-Resolution Mass Spectrometry of purified compounds

High Resolution Mass Spectrometry was acquired on a C18 column (Thermo Acclaim 120 C_18_, 2.1 × 150 mM) using a Dionex U-3000 HPLC system connected to an LTQ-Orbitrap Mass Spectrometer (Thermo-Fisher).

### Murine work

All experimental procedures were approved by the Animal Care and Use Committee of The Icahn School of Medicine at Mount Sinai (PI Cohen IACUC-2016-0491). Germ free C57BL/6 mice were maintained in sterile isolators with autoclaved food and water in the Gnotobiotic Facility of the Faith Lab at Mount Sinai. 6-week-old mice were used for all experiments (3M and 3F in the treatment group, 5M and 1F in the control group). The treatment group was colonized with the SIHUMI whereas the control group was left germ free. For colonization studies 5 ml of an overnight culture in LBM media of the SIHUMI (treatment group) was centrifuged at 500 x g for 2 minutes, the supernatant was decanted and the cells were resuspended in 2 ml of sterile PBS. Germ free mice were gavaged with 100 µL of bacterial culture immediately upon removal from sterile isolators. Colonization was confirmed by collection of fecal pellets after 3 days. Crude DNA was extracted from fecal pellets per protocol (ZymoBIOMICS DNA/RNA Miniprep Kit) and colonization conformed by targeted PCR of each strain using specific primers as detailed above. After colonization mice were housed in specific-pathogen-free conditions and fed with autoclaved water and food. After colonization for 7 days the mice were euthanized and samples were collected for analysis. 200 mg of cecal contents were collected from each mouse and placed immediately at −80 °C. The animal experiments were not randomized and the investigators were not blinded to the allocation during experiments and outcome assessment. No statistical methods were used to predetermine sample size. All mice which completed the experiments were analyzed.

### Metabolite quantitation by mass spectrometry

Cecal samples were weighed into 2 mL microtubes containing 2.8 mm ceramic beads (Omni International) and resuspended to a final concentration of 100 mg/mL using 80:20 methanol:water containing phenol-d7, palmitic acid-d31 and ^13^C,^15^N-amino acid internal standards (Cambridge Isotope Laboratories). Homogenization was using a Bead Ruptor (Omni International) at 6 m/s for 30 s for 6 cycles, at 4 °C. Samples were centrifuged for 20 minutes at 20,000 × g at 4 °C and then divided for 3 analytical methods.

### Method 1: GC-nCI-MS with PFB Derivatization

100 µL of cecal extract was added to 100 µL of 100 mM borate Buffer (pH 10), 400 µL of 100 mM pentafluorobenzyl bromide (Thermo Scientific) in acetone (Fisher), and 400 µL of cyclohexane (Acros Organics) in a sealed autosampler vial. Samples were heated to 65 °C for 1 hour with shaking. After cooling to room temperature and allowing the layers to separate, 100 μL of the cyclohexane upper phase was transferred to autosampler vial containing a glass insert and sealed. Analyzed was using an GC-MS (Agilent 7890A GC system, Agilent 5975C MS detector) operating in negative chemical ionization mode, using a DB-5MS column (30 m x 0.25 mm, 0.25, 0.25 μm; Agilent Technologies), methane as the reagent gas and 1 μL split injection (1:5 split ratio). Raw peak areas for aromatic analytes (tyramine and hydrocinnamic acid) were normalized to phenol-d7 internal standard and lipid analytes (9,10-methylenehexadecanoic acid and 12-methyltetradecanoic acid) were normalized to palmitic acid-d31 internal standard. Data analysis was performed using MassHunter Quantitative Analysis software (version B.09, Agilent Technologies).

### Method 2: LC triple quadrupole with reverse phase chromatography

200 µL of extract was dried using a vacuum concentrator (Genevac) and resuspended in 400 µL 50:50 methanol:water, clarified by centrifugation and analyzed suing reverse phase chromatography coupled to TSQ Vantage triple quadupole mass spectrometer with HESI II source. LC separation was using an HSS T3 column (100 × 2.1 mm, 1.8 μm particle size, Waters) and Agilent 1260 binary pump. Mobile phase A was 0.1% formic acid in water and Mobile phase B was 0.1% formic acid in acetonitrile. The gradient was 0 min, 0% B; 2 min, 0% B; 5 min, 12% B; 7 min, 70% B; 8.5 min, 97% B, 11.5 min, 97% B with 3.5 min of re-equilibration time. LC parameters were: flow rate 300 μL/min, injection volume 15 μ and column temperature 35 °C. The mass spectrometer was operated in positive ionization with transitions for tyrptamine (m/z 161.1 → 115.1, CE 30V*; 161.1 → 144.1, CE 4 V) and nicotinic acid (m/z 124.1 → 80.1, CE 18 V*; 124.1 → 78.1, CE 19V), with * indicating the primary transition used for quantitation. MS parameters were: capillary temp: 300°C; vaporizer temp: 350 °C; sheath gas: 50; aux gas: 30; spray voltage 4000 V. Data was acquired and analyzed using TraceFinder software (version 4.1, Thermo Scientific) confirmed by comparison with authentic standards.

### Method 3: LC Q-TOF with HILIC chromatography

Samples were prepared as for Method 2 but then resuspended in 200 μL 60:40 acetonitrile:water and analyzed by hydrophilic interaction chromatography (HILIC) coupled to the 6545 Q-TOF mass spectrometer with Dual JetStream source (Agilent). The LC separation was using an Acquity UPLC BEH Amide column (150 mm × 2.1 mm, 1.7 μm particle size, Waters) and Agilent 1290 Infinity II binary pump. Mobile phase A was 90:10 water:acetonitrile with 10 mM ammonium acetate and 0.2% acetic acid, and mobile phase B was 10:90 wate:acetonitrile with 10 mM ammonium acetate and 0.2% acetic acid. The gradient was 0 min, 95% B; 9 min, 70% B; 10 min, 40% B; 13 min, 30% B; 15 min, 95% B. LC parameters were: flow rate 400 μL/min, column temperature 40 °C, and injection volume 5 μL. The mass spectrometer was operated in positive ionization mode. MS parameters were: gas temp: 325 °C; gas flow: 10 L/min; nebulizer pressure: 35 psig; sheath gas temp: 400 °C; sheath gas flow: 12 L/min; VCap: 4,000 V; fragmentor: 125 V. Active reference mass correction was done through a second nebulizer using masses with m/z: 12.050873 and 922.009798. Data were acquired over m/z range 50–1700 and analyzed using MassHunter Profinder software (version B.09, Agilent) and confirmed by comparison with a cadaverine authentic standard. Compiling these data sets in GraphPad Prizm was then used to derive p-values. Unpaired t test (two-tailed) were used.

### Tyramine

Fraction 3 from *E. coli* LF82 was chosen as the pilot fraction for the dopamine receptors. 1 mL of Fraction 3 (100 mg/mL in DMSO) was dried down resuspended in 1 mL 50/50 MeOH:H2O and injected in 50 µL increments onto a semi-preparative 250 × 10 mm Luna^®^ Omega 2.6 uM Polar C18 LC column on an Agilent 1100 HPLC with a solvent system where Solvent A was H_2_O + 0.1% formic acid and Solvent B was CH_3_CN + 0.1% formic acid. The chromatographic method was as follows: 0% B for 5 CV, then up to 90% B over 15 CV, with a 5 CV hold at 90% B. Peak detection and fraction collection was driven by UV absorbance at 210 nm, 254 nm, 280 nm, and 330 nm. Fractions were collected and re-assayed against DRD3 to guide further purification. The active fraction was further purified using a 150 x 10 mm Kinetix^®^ 5 μm Biphenyl 100A LC column. A single resulting fraction retained activity and this compounds was identified as tyramine by NMR and HRMS (LC-HRMS-ESI (*m/z*): [M+H]^+^ calcd for C_8_H_11_NO, 138.0841; found 138.0911). Tyramine: ^1^H NMR (DMSO-*d*_6_, 600 MHz): δ_H_ 7.02 (2H, d, *J* = 8.5 Hz), 6.70 (2H, d, J = 8.5 Hz), 2.89(2, t, J = 7.6 Hz), 2.71 (2H, t, J = 7.6 Hz). ^13^C NMR (DMSO-*d*_6_, 151 MHz): δ_C_ 156.1 (1C, s), 129.5 (2C, s), 115.3 (2C, s), 40.8 (1C, s), 33.45 (1C, s).

### Tryptamine

A single fraction from the fermentation of *Ruminococcus gnavus* was chosen as a pilot fraction to find serotonin active compounds which could then be assessed in other bacteria. 1 mL of Fraction 5 solution (100 mg/mL in DMSO) was dried down, resuspended in 1 mL 50/50 MeOH:H_2_O, and injected in 50 µL increments onto a semi-preparative 250×10 mm Luna^®^ Omega 2.6 µM Polar C18 LC column on an Agilent 1100 HPLC with a solvent system where Solvent A was H2O + 0.1% formic acid and Solvent B was CH3CN + 0.1% formic acid. The chromatographic method was as follows: 0% B for 5 CV, then up to 90% B over 15 CV, with a 5 CV hold at 90% B. Peak detection and fraction collection was driven by UV absorbance at 210 nm, 254 nm, 280 nm, and 330 nm. Fractions were collected and re-assayed against HTR5A to guide further purification. The active fraction (41 mg) was ~90% tryptamine as evident by NMR and HRMS (LC-HRMS-ESI (*m/z*): [M+H]^+^ calcd for C_8_H_11_NO, 161.1000; found 161.1071). Tryptamine: ^1^H NMR (DMSO-*d*_6_, 600 MHz): δ_H_ 7.54 (1H, d, *J* = 7.9 Hz), 7.20 (1H, d, 2.0 Hz), 7.08 (1H, t, 7.6 Hz), 7.00 (1H, t, 7.6 Hz), 3.01 (2H, dd, 8.5 Hz, 7.1 Hz), 2.93 (2H, dd, 8.8, 6.2 Hz). ^13^C NMR (DMSO-*d*_6_, 151 MHz): δ_C_ 136.3 (1C, s), 126.9 (1C, s), 123.2 (1C, s), 121.1 (1C, s), 118.4 (1C, s), 118.1 (1C, s), 111.5 (1C, s), 110.1 (1C, s), 40.0 (1C, s), 24.6 (1C, s).

### Phenylpropanoic acid

Due to its relative simplicity in composition, Fraction 2 was chosen for further study. 40 mg of Fraction 2 was injected in two equal increments onto a semi-preparative 150 × 10 mm XBridge^®^ 5 μm C18 columnon an Agilent 1100 HPLC with a solvent system where Solvent A was H_2_O + 0.1% formic acid and Solvent B was CH_3_CN + 0.1% formic acid. The chromatographic method was as follows: flow rate 4 mL/min; 2.5% B for 5 min, then increased to 35% B over 25 min, then flushed at 99% B for 5 min. Peak detection and fraction collection was driven by UV absorbance at 210 nm, 254 nm, 280 nm, and 330 nm. Fractions were collected per minute and re-assayed against GPR109B. All activity was found in subfractions 30 (8.8 mg) and 31 (0.1 mg), which were identified as pure phenylpropanoic acid by NMR and MS. Significant quantities of phenylpropanoic acid was also subsequently detected in Fraction 7 from all bacterial extracts. UPLC-MS-ESI (*m/z*): [M]^-^calcd for C_9_H_9_O_2_149.06, found 149.03.^1^H NMR (DMSO-*d*_6_, 600 MHz): δH 12.17 (1H, bs), 7.27 (2H, t, J = 6.9 Hz), 7.22 (2H, d, J = 7.3 Hz), 7.18 (1H, t, J = 7.0 Hz), 2.81 (2H, t, J = 7.8 Hz), 2.51 (2H, t, J = 7.9 Hz). ^13^C NMR (DMSO-d6, 151 MHz): δC 174.0 (1C, s), 141.0 (1C, s), 128.3 (2C, s), 128.2 (2C, s), 125.9 (1C, s), 35.5 (1C, s), 30.5 (1C, s).

### Cadaverine

Fraction 4 from *E. coli* LF82 was chosen as the pilot fraction for the histamine receptors. 1 mL of Fraction 4 (100 mg/mL in DMSO) was dried down resuspended in 1 mL H2O and injected in 50 µL increments onto a semi-preparative 250 x 10 mm Luna^®^ Omega 2.6 uM Polar C18 LC column on an Agilent 1100 HPLC with a solvent system where Solvent A was H2O + 0.1% formic acid and Solvent B was CH3CN + 0.1% formic acid. The chromatographic method was as follows: 0% B for 10 CV, then up to 90% B over 5 CV, with a 3 CV hold at 90% B. Peak detection and fraction collection was driven by charged aerosol detection using a Corona Veo (ThermoFisher Scientific) after UV proved to not be useful. Fractions were collected and re-assayed against HRH4 to guide further purification. The active fraction was further purified two more times using the same Polar C18 column with extended flushes at 0% B, as the activity always was eluting in the void. A HILIC method proved to be less effective. A single resulting fraction retained activity and this compound was identified as cadaverine by NMR and HRMS (LC-HRMS-ESI (*m/z*): [M+H]^+^ calcd for C_5_H_14_N_2_, 102.1157; found 102.12293). Co-eluted in this fraction was the compound agmatine (LC-HRMS-ESI (*m/z*): [M+H]^+^ calcd for C_5_H_14_N_4_, 130.12184; found 131.12920). Cadaverine: ^1^H NMR (D_2_O, 600 MHz): δ_H_ 3.04 (4H, t, *J* = 7.6 Hz), 1.74 (4H, p, *J* = 7.7 Hz), 1.49 (2H, p, *J* = 7.7 Hz). ^13^C NMR (D_2_O, 151 MHz): δ_C_ 39.2 (2C, s), 26.2 (2C, s), 22.7 (1C, s).

### 9,10-methylenehexadecanoic acid

Fraction 9 of *E. coli* LF-82 was injected in DMSO onto a semi-preparative 150 × 10 mm XBridge^®^ 5 μm C18 column with a solvent system where Solvent A was H2O + 0.1% formic acid and Solvent B was CH3CN + 0.1% formic acid. The chromatographic method was as follows: 30% B for 3 column CV then up to 99% B over 5 CV, with a 15 CV hold at 99% B. Peak detection and fraction collection was driven by charged aerosol detection using a Corona Veo (ThermoFisher Scientific) after UV proved to not be useful. Fractions were collected and re-assayed against BAI1 to guide further purification. A single resulting fraction retained activity and this compound was identified as 9,10-methylenehexadecanoic acid by NMR and HRMS (LC-HRMS-ESI (*m/z*): [M+H]^+^ calcd for C_17_H_32_O_2_, 267.2402; found 267.2334). 9,10-methylenehexadecanoic acid: ^1^H NMR (CDCl_3_, 600 MHz): δ_H_ 2.35 (2H, t, *J* = 7.4 Hz), 1.64 (2H, p, *J* = 7.4 Hz), 1.37 (16H, m), 1.32 (2H, m), 1.14 (2H, m), 0.89 (3H, t, *J* = 6.6 Hz), 0.65 (2H, m), 0.57 (1H, td, J = 8.2 Hz, 4.2 Hz), −0.33 (1H, q, J = 5.2, 4.4 Hz). ^13^C NMR (CDCl_3_, 151 MHz): δC 177.7 (1C, s), 33.8 (1C, s), 32.2 (1C, s), 30.4 (1C, s), 30.4 (1C, s), 29.7 (1C, s), 29.6 (1C, s), 29.5 (1C, s), 29.3 (1C, s), 29.0 (1C, s), 28.9 (1C, s), 25.0 (1C, s), 23.0 (1C, s), 16.0 (1C, s), 16.0 (1C, s), 14.4 (1C, s), 11.2 (1C, s).

### 12-methylmyristic acid

Fraction 9 of *B. vulgatus* was injected in DMSO onto a semi-preparative 150 × 10 mm XBridge^®^ 5 μm C18 column with a solvent system where Solvent A was H2O + 0.1% formic acid and Solvent B was CH3CN + 0.1% formic acid. The chromatographic method was as follows: 30% B for 3 column CV then up to 99% B over 5 CV, with a 15 CV hold at 99% B. Peak detection and fraction collection was driven by charged aerosol detection using a Corona Veo (ThermoFisher Scientific) after UV proved to not be useful. Fractions were collected and re-assayed against BAI1 to guide further purification. A single resulting fraction retained activity and this compound was identified as 9,10-methylenehexadecanoic acid by NMR and HRMS. (LC-HRMS-ESI (*m/z*): [M-H]^−^ calcd for C_15_H_30_O_2_, 241.2245; found 241.2178). 12-methylmyristic acid: ^1^H NMR (CDCl_3_, 600 MHz): δH 2.35 (2H, t, 7.5 Hz), 1.64 (2H, p, 7.5 Hz), 1.26 (16H, m), 1.12 (1H, m, 6.9 Hz), 1.08 (2H, m), 0.85 (3H, t, 7.4 Hz), 0.84 (3H, d, 5.1 Hz). ^13^C NMR (CDCl3, 151 MHz): δC 178.2 (1C, s), 36.6 (1C, s), 34.3 (1C, s), 33.7 (1C, s), 30.0 (1C, s), 29.6 (1C, s), 29.5 (1C, s), 29.4 (1C, s), 29.4 (1C, s), 29.2 (1C, s), 29.0 (1C, s), 27.0 (1C, s), 24.6 (1C, s), 19.2 (1C, s), 11.4 (1C, s).

**Figure S1.**
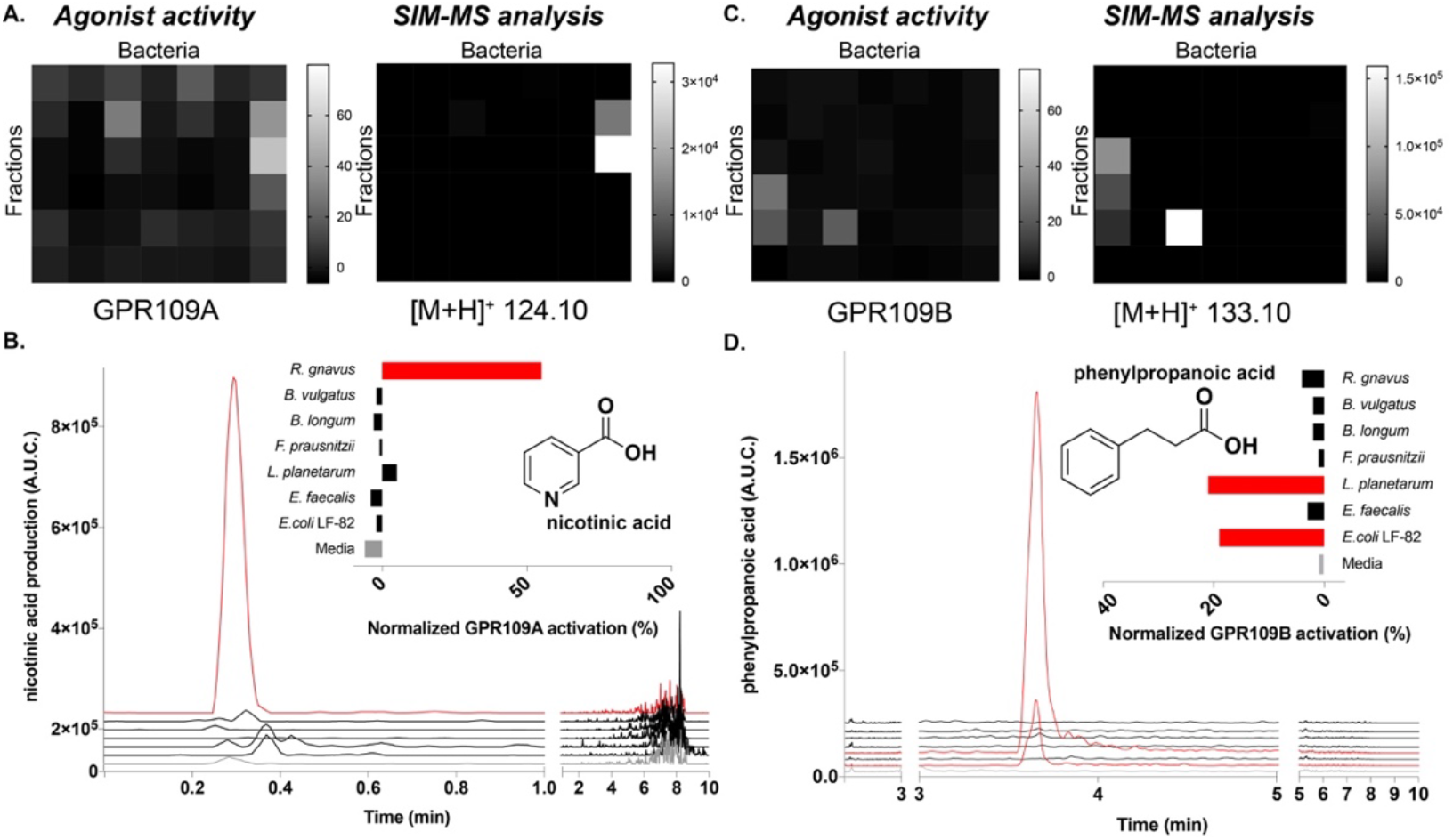
Heatmaps of GPR109A agonism activity and MS-based detection of nicotinic acid in early, polar fractions 1-6. A. Heatmaps of GPR109A agonism activity and MS-based detection of nicotinic acid in early, polar fractions 1-6. B. GPR109A agonism and SIM-MS analysis of nicotinic acid of fraction 3 across all SIHUMI members. C. Heatmaps of GPR109B agonism activity and MS-based detection of phenylpropanoic acid in early, polar fractions 1-6. D. GPR109B agonism and SIM-MS analysis of phenylpropanoic acid of fraction 3 across all SIHUMI members.

**Figure S2.**
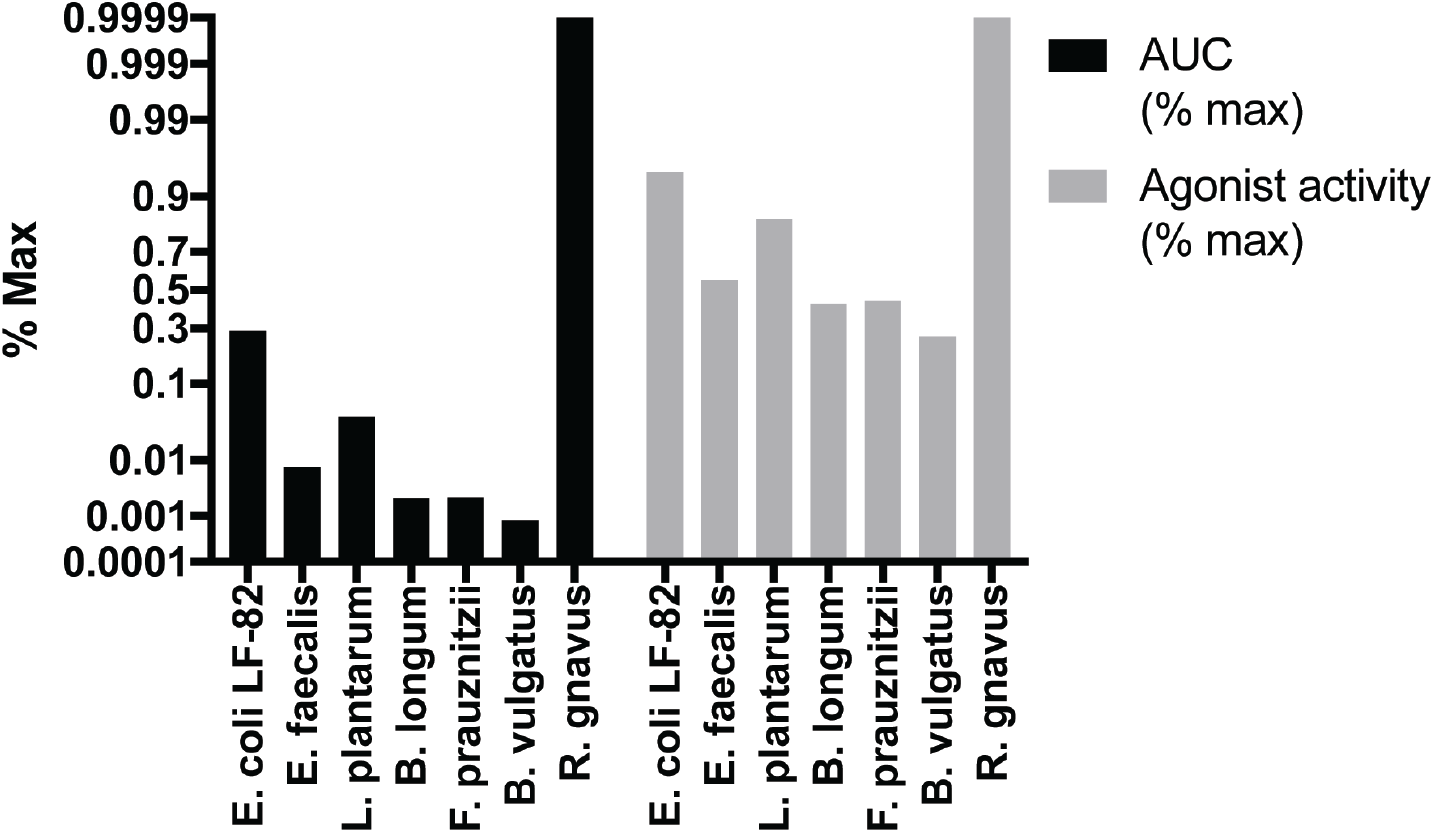
Correlation of chemical MS-based identification of tyramine and dopamine receptor activation in representative fraction from each bacterium. Correlation of chemical MS-based identification of tyramine and dopamine receptor activation in representative fraction from each bacterium. Values represent peak height of ion extractions for the mass of tyramine and the activation of the dopamine receptor DRD3 normalized to maximum peak height and agonist activity, respectively.

**Figure S3.**
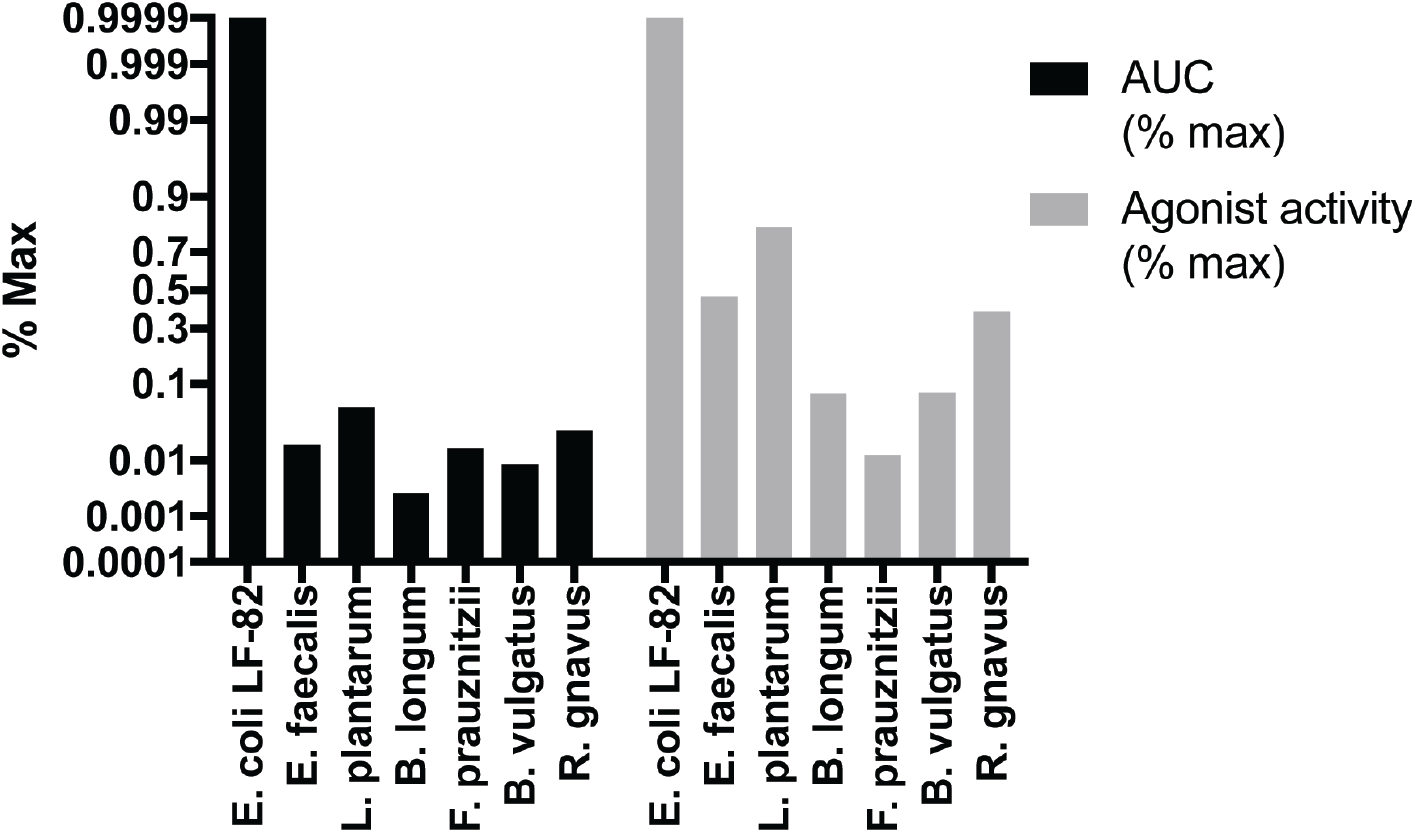
Correlation of chemical MS-based identification of tryptamine and serotonin receptor activation in representative fraction from each bacterium. Correlation of chemical MS-based identification of tryptamine and serotonin receptor activation in representative fraction from each bacterium. Values represent peak height of ion extractions for the mass of tryptamine and the activation of the serotonin receptor HTR2A normalized to maximum peak height and agonist activity, respectively.

**Figure S4.**
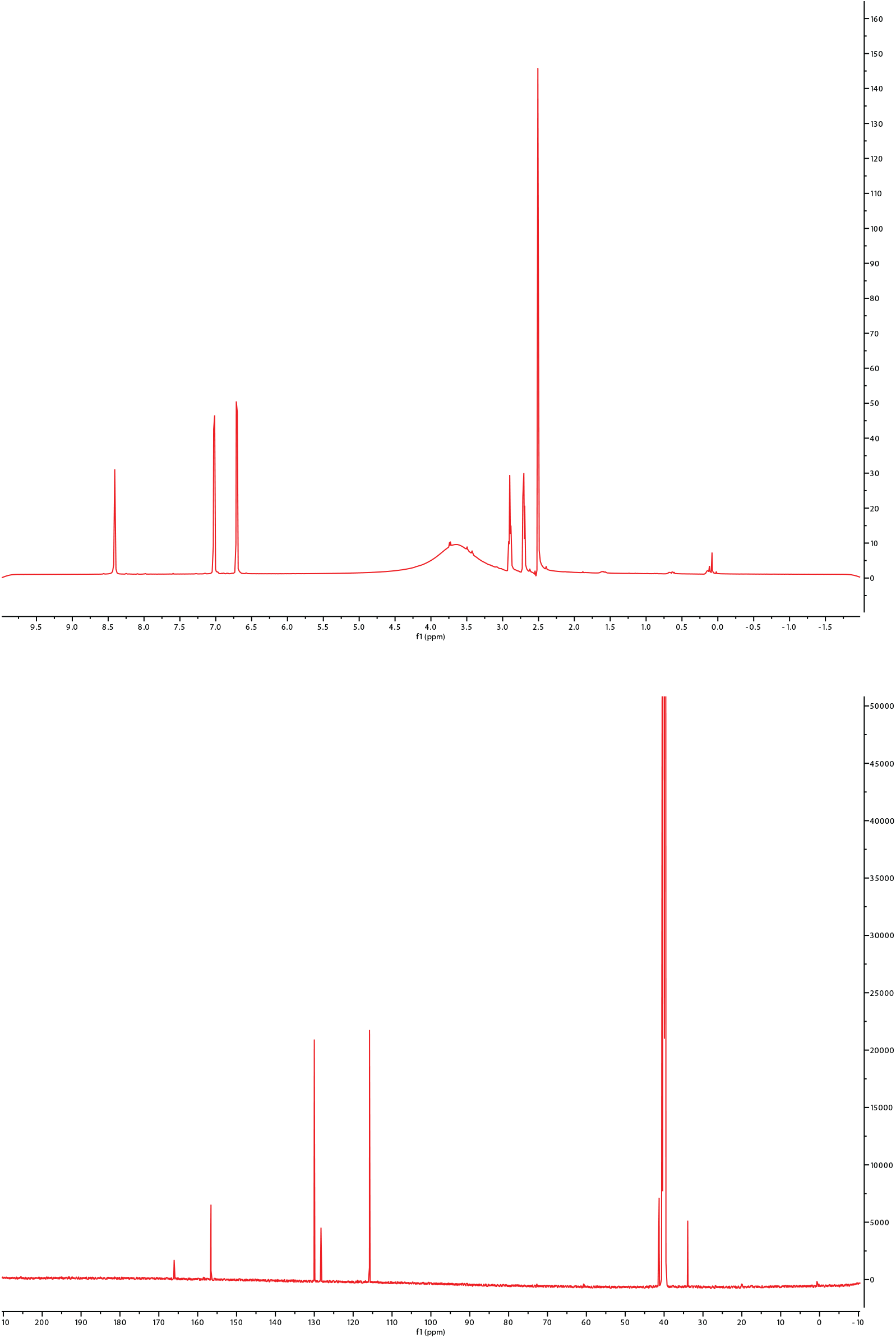
^1^H and ^13^C NMR data for isolated tryptamine (DMSO-*d*_6_).

**Figure S5.**
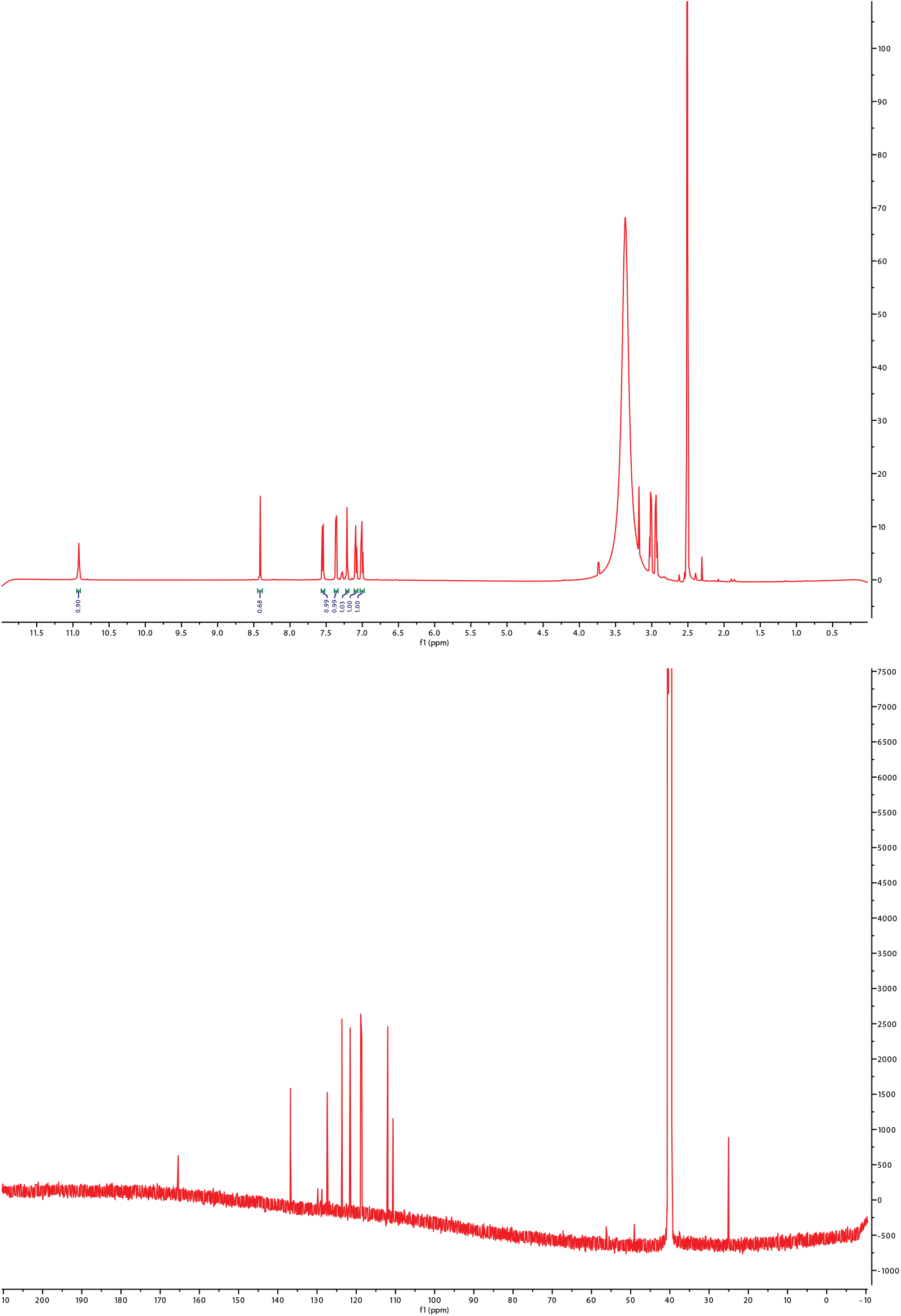
^1^H and ^13^C NMR data for isolated tryptamine (DMSO-*d*_6_).

**Figure S6.**
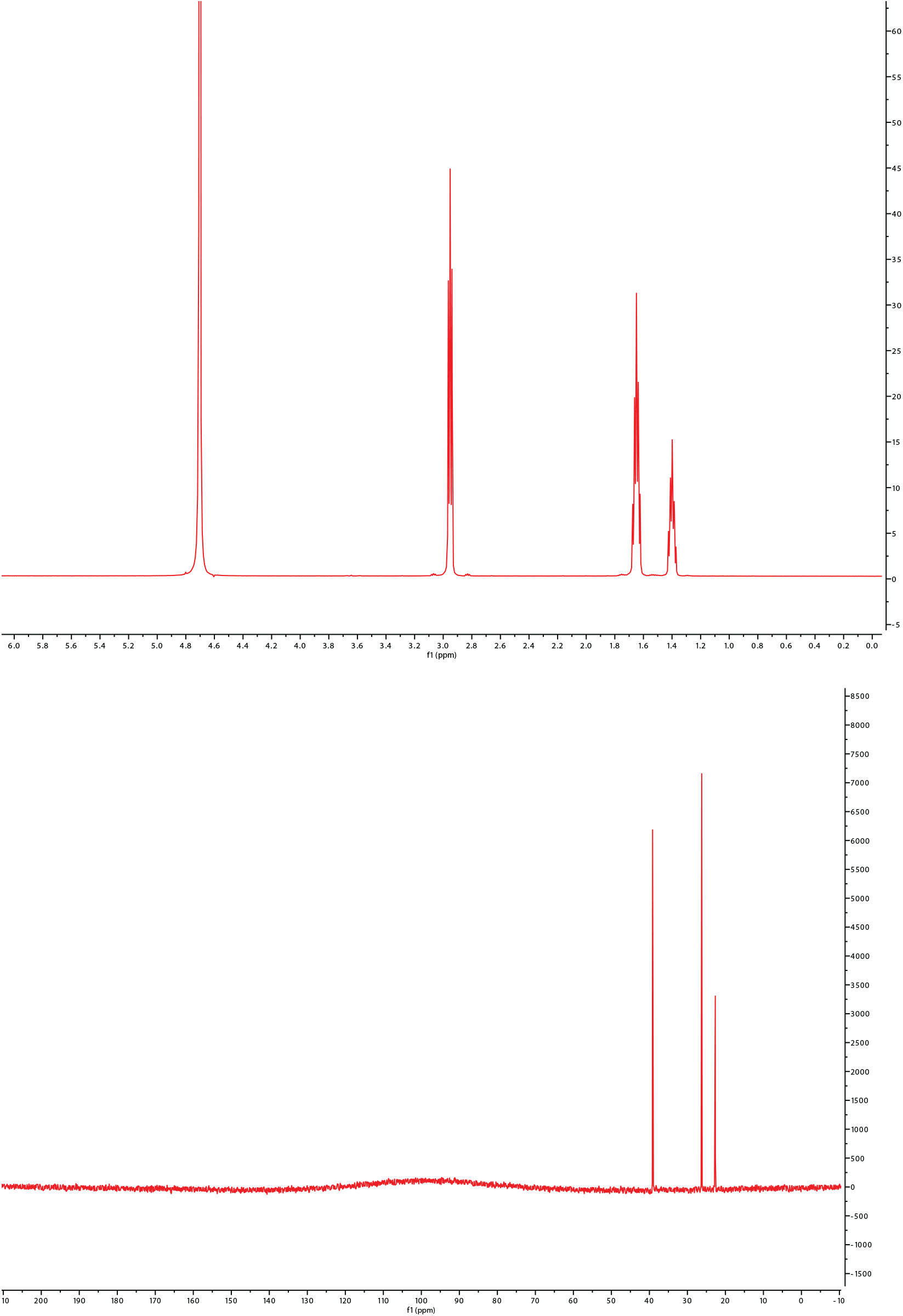
^1^H and ^13^C NMR data for isolated cadaverine (D_2_O)

**Figure S7.**
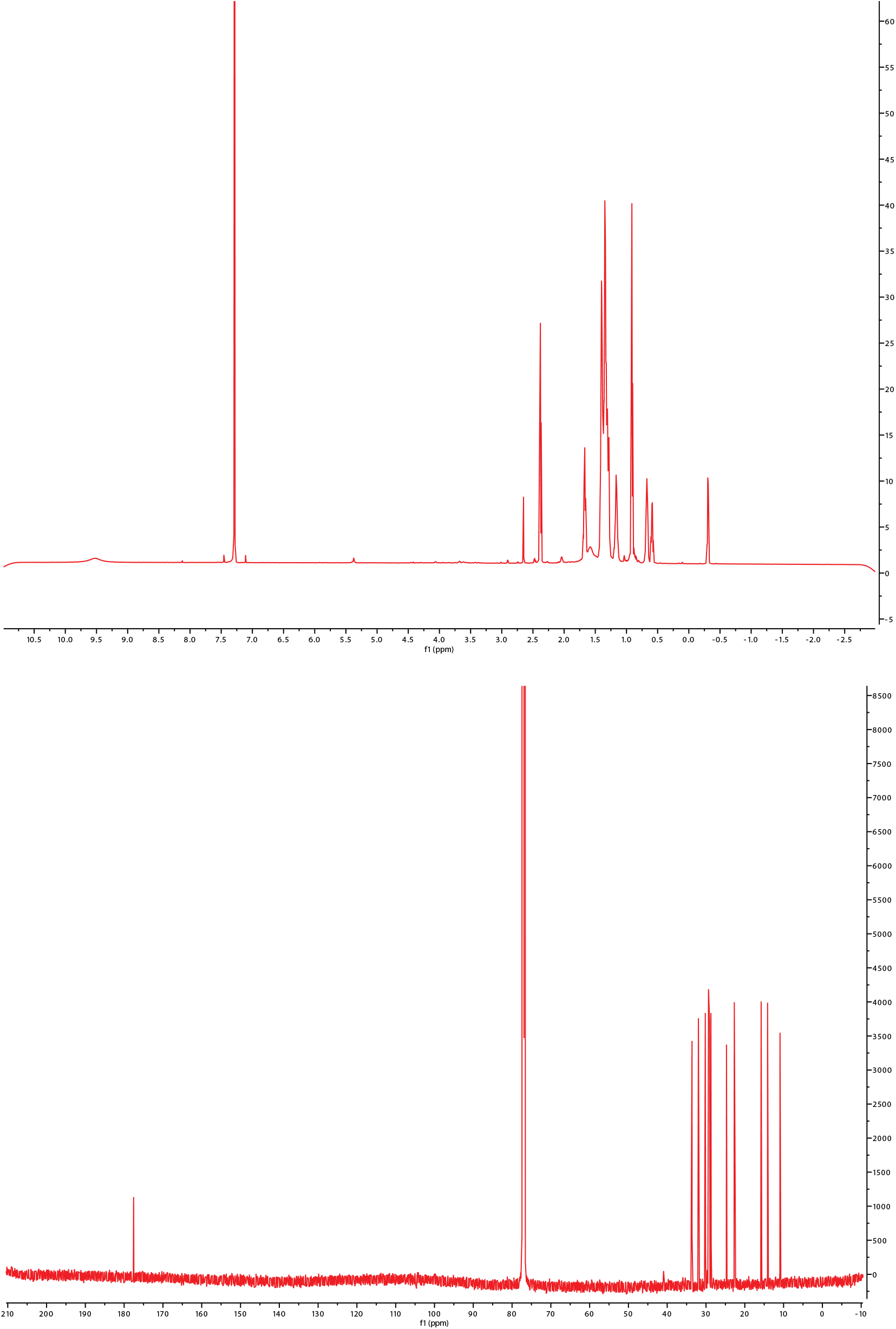
^1^H and ^13^C NMR data for isolated 9,10-methylenehexadecanoic acid (CDCl_3_)

**Figure S8.**
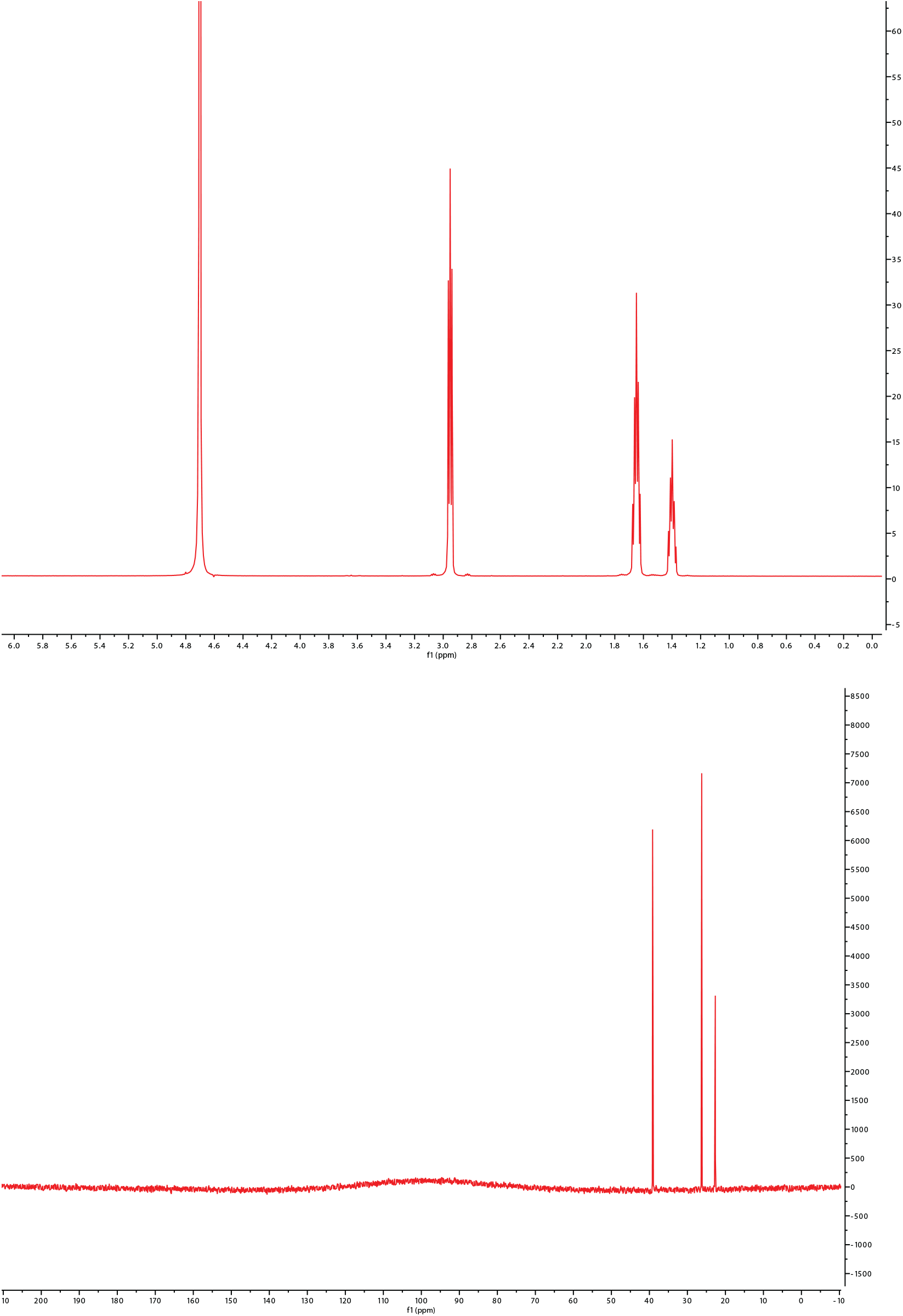
^1^H and ^13^C NMR data for isolated 12-methyltetradecannoic acid (CDCl_3_)

**Figure S9.**
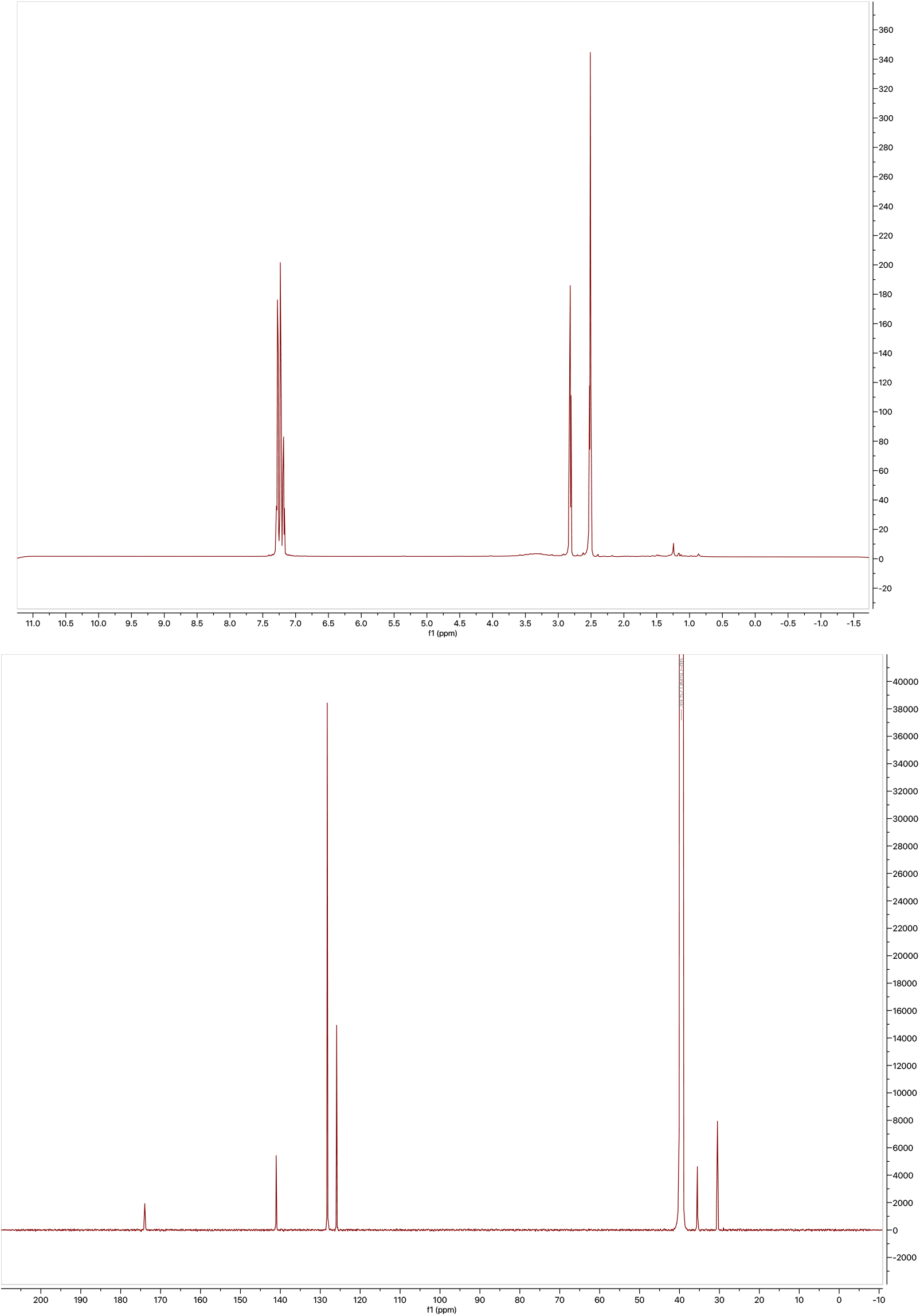
^1^H and ^13^C NMR data for isolated phenylpropanoic acid (DMSO-d_6_)

## Additional data table S1 (separate file)

Table S1 List of GPCRs tested with control compounds indicated

Table S2 Raw screening data for GPCR agonist screen with tested positive controls

Table S3 Raw screening data for Orphan GPCR agonist screen

Table S4 List of hit filtering by receptor type

Table S5 a. Raw cadA BLAST analysis of HMRGD. B. cadA PFAM analysis of HMRGD

Table S7 Raw targeted HRMS data

